# Falls Induced by Optogenetic Inhibition of Basal Forebrain Cholinergic Projections after Dorsomedial Striatal Dopamine Depletion in a Dual Disruption Model of Falling Vulnerability in Parkinson Disease

**DOI:** 10.64898/2026.06.04.729642

**Authors:** Aaron Kucinski

## Abstract

Falls are a common and debilitating feature of Parkinson’s Disease (PD) patients. Prefrontal acetylcholine (ACh) deficits, as well as nigrostriatal dopamine deficits, are implicated in vulnerability to falls. PD patients with loss of cortical ACh and associated cognitive dysfunction experience a higher rate of falls than PD patients without cortical ACh loss. In addition, chemogenetic inhibition of basal forebrain (BF) neurons in rats increases the vulnerability to falls on a balance beam task. Here, the impact of transient optogenetic inhibition specifically of BF cholinergic neurons was assessed in rats with dorsomedial striatal dopamine lesions during traversal of straight or zig-zag balance beams using the Michigan Complex Movement Control Task (MCMCT). Adding transient optogenetic inhibition of BF cholinergic neurons with striatal dopamine lesions elevated falls above the level produced by striatal dopamine lesions or BF ACh inhibition alone, especially on the challenging zig-zag task. These results support the critical role of BF-cortical cholinergic circuits in alleviating vulnerability to falls in PD patients with striatal dopamine loss, suggesting that it is combined loss of BF cholinergic activity and striatal dopamine that leads to greatest vulnerability to falls and related complex movement impairments.

## Introduction

Parkinson’s Disease (PD) is a multi-system neurodegenerative disorder that involves changes in multiple neurotransmitter systems. Loss of dopamine (DA) in the dorsal striatum is the principal neurotransmitter deficit responsible for the characteristic PD movement impairments. Changes in cholinergic systems also play key roles in disease pathophysiology (Bohnen et al., 2022). PD patients with regional cholinergic deficits often develop additional features including cognitive deficiencies (Bohnen et al., 2006; Schliebs & Arendt, 2011), as well as levodopa-resistant movement deficits such as impaired gait, poor balance and posture, and a higher propensity for falls than PD patients without cholinergic losses (Balash et al., 2005; Rochester et al., 2012). Falls are dangerous and a leading cause of death in PD patients and in the elderly in general, particularly in those with cognitive decline (Balash et al., 2005; Grimbergen et al., 2004; Holtzer et al., 2007). PD patients that fall tend to do so recurrently, especially patients with brain cholinergic deficits (Bohnen et al., 2009) and those with cognitive dysfunction such as poor attention (Allcock et al., 2009; Yarnall et al., 2011).

Cholinergic output of the basal forebrain (BF), especially projections to prefrontal cortex, regulates cognitive processes (Gombkoto et al., 2021; Liu et al., 2015; Sarter & Paolone, 2011; Sarter & Lustig, 2020; Záborszky et al., 2018) including the detection and integration of cues necessary to guide complex movement (Albin et al., 2022; Gritton et al., 2016). Degeneration of BF-cortical cholinergic pathways contributes to declining cognition in humans (Ballinger et al., 2016; Düzel et al., 2010) and rodents (Sarter, 1998). Loss of BF cholinergic neurons and reduced cortical ACh has also been described in PD patients with impaired cognition (d’Angremont et al., 2025; Grothe et al., 2021; Schulz et al., 2018; Wolf et al., 2014), including poor attention (Woollacott & Shumway-Cook, 2002), and in patients with gait impairments such as high fall propensity (Dalrymple et al., 2021; LaPointe et al., 2010), particularly while traversing complex surfaces or dual-tasking (Kim et al., 2019).

A rat model of high fall propensity was previously developed using dual-system lesions targeting both the DA and ACh neurotransmitter systems affected in PD (Kucinski et al., 2013; Sarter et al., 2014). Rats with partial losses of dorsal striatal DA terminals following striatal 6-OHDA infusions combined with loss of cortical ACh after BF lesions exhibited heightened propensity for falls on a challenging balance-beam traversal task, the Michigan Complex Movement Control Task (MCMCT), as well as reduced attentional performance on a sustained attention task (SAT). Decreasing cholinergic output from BF to the cortex was also achieved using reversible chemogenetic deactivation of ACh systems with Designer Receptors Exclusively Activated by Designer Drugs (DREADDS) to transiently inhibit BF neurons and thus induce falls, especially in rats with more cognitively guided motor control (Kucinski et al., 2019; Kucinski et al., 2022).

The current study sought to replicate and extend previous findings of elevated fall propensity with BF inhibition, particularly together with DA losses, by combining striatal 6-OHDA lesions with selective cholinergic BF neuronal inhibition using wireless LED optogenetics to transiently disrupt BF cholinergic activity on a sub-second time scale. Vulnerability to falls was assessed using the MCMCT, as rats traversed either a rotating straight balance beam or a more challenging rotating beam shaped in a zig-zag pattern.

Previous research found that disruption of ACh activity induced by nonspecific chemogenetic inhibition of BF neurons was more likely to increase falls by goal tracking rats (Goal Trackers; GT) than by sign tracking rats (Sign Trackers; ST) (Kucinski et al., 2019). Sign-tracking is assessed in Pavlovian reward conditioning tests as a tendency to approach and engage with reward-associated cues (CS+) and is suggested to be associated with relatively greater vulnerability for developing and maintaining addiction-like behaviors due to increased responsiveness to salient stimuli (Pitchers et al., 2017a; Pitchers et al., 2017b; Robinson et al., 2014; Yager & Robinson, 2013). Goal-tracking, on the other hand, is classified as approach directly to the location of rewards themselves, and is associated with heightened release of cortical ACh during a sustained attention task (Paolone et al., 2013), elevated presynaptic choline uptake in BF cholinergic neurons (Koshy Cherian et al., 2017), and a more top-down, cognitive strategy on complex beam traversal tasks relative to STs (Kucinski et al., 2018). The fall propensities of GT and ST were further compared here using the MCMCT while testing the effects of dual ACh and DA disruptions produced by optogenetic cholinergic inhibition of the BF together with depletion of DA in the dorsal striatum.

## Methods

### Subjects and Housing

ChAT-Cre heterozygous male and female Long–Evans rats between 2 and 3 months of age were obtained from Stanford University (Department of Bioengineering) and bred with wild-type (WT) Long–Evans female rats obtained from Envigo, and genotyped by Transnetyx using tail snips. Animals were individually housed in opaque single standard cages (27.70 cm × 20.30 cm) in a temperature- and humidity-controlled environment (23 °C, 45%) and maintained under a 12:12 h light/dark schedule (lights on at 7:00 AM). Food (Envigo Teklad rodent diet) and water were available *ad libitum*. All procedures were conducted in adherence with protocols approved by the University Committee on Use and Care of Animals at the University of Michigan and in laboratories accredited by the Association for Assessment and Accreditation of Laboratory Animal Care.

### Sign and Goal Trackers

#### GT/ST Screening

Following genotyping, ChAT-Cre positive rats underwent Pavlovian Conditioned Approach (PCA) screening for individual propensity toward goal-tracking versus sign-tracking (Meyer et al., 2012). PCA screening of 163 rats in total yielded 41 GTs (17 females) and 56 ST (23 females) (see Table 1). 27 GTs (15 females) and 30 STs (20 females) were used to assess the effects of optogenetic deactivation of cholinergic neurons of the BF on MCMCT performance measures. Rats were aged 3-4 months during PCA screening and 4-6 months during MCMCT testing. Following completion of the MCMCT test battery, rats were perfused for histological analyses.

**Table 1.**
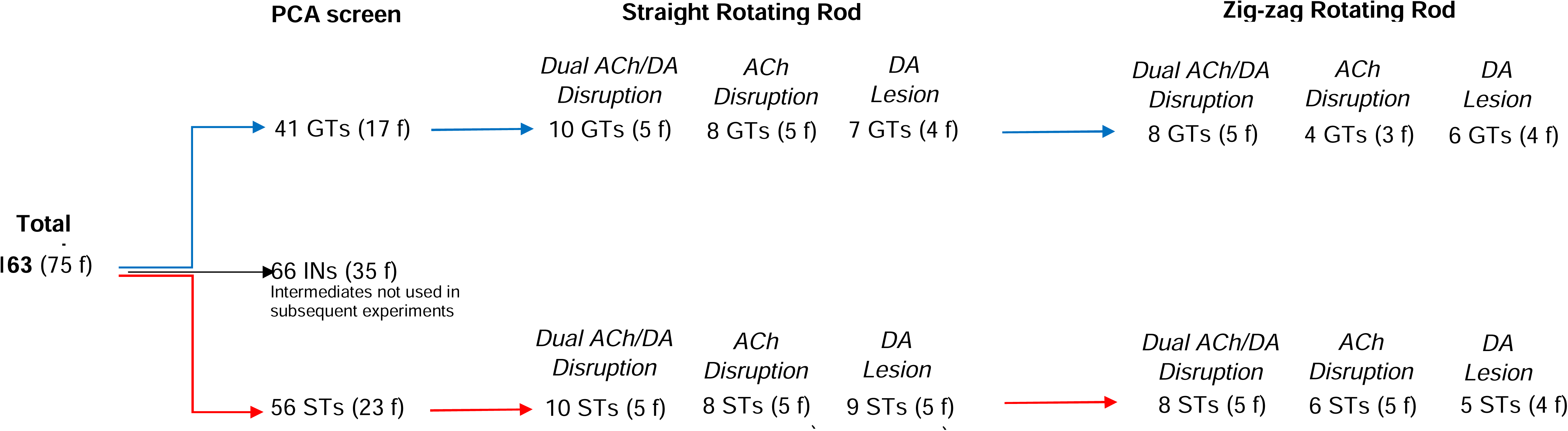
Number of Chat-Cre positive rats undergoing PCA screening and Goal Trackers (GT) and Sign Trackers (ST) in each Disruption Group used to assess MCMCT falls from the straight and zig-zag rods with continuous LED inhibition (f, females).

### PCA Screening for STs vs GTs

#### Apparatus and procedures

The screening of STs and GTs using a PCA test followed established and previously described methods (e.g., Paolone et al., 2013; Pitchers et al., 2017a; Yager et al., 2015). Briefly, rats were handled daily for 3 days and given ∼15 banana-flavored grain-based pellets (45 mg; BioServ) in their home cages for 2 d prior to the start of PCA testing. Rats were tested in conditioning chambers (20.5 × 24.1 cm floor area, 20.2 cm high; MED Associates Inc.). Throughout the duration of the experiments, males and females were tested in separate chambers. A food magazine in the center of one of the walls of the chamber with an automatic feeder delivered banana pellets. Infrared photobeam breaks detected magazine entries. On either side of the magazine was a retractable lever with an LED backlight illuminated only when the lever extended into the chamber. Deflections of the lever were used to quantify lever contacts. The beginning of a test session was signaled by the illumination of a red house light located near the ceiling of the side of the chamber opposite to the magazine/lever. On the first day of testing (“pretraining”), rats were placed into the conditioning chambers and the house light was illuminated after a 5 min habituation period. Twenty-five pellets were then delivered on a VI-30 (0–60 s) schedule. On average, this pretraining session lasted 12.5 min and the lever was retracted throughout the session. During this session and all subsequent PCA sessions, rats consumed all pellets. The house light was turned on in the next five PCA sessions and rats were presented with 25 lever/pellet pairings delivered on a VI-90 (30–150 s) schedule. The CS for each trial was the extension of the illuminated lever into the chamber for 8 s. After lever retraction, a pellet was immediately delivered into the magazine. On average, the PCA test sessions lasted 37.5 min.

#### PCA measures and classification criteria

Lever presses and magazine entries during the CS periods were used to quantify three measures of approach that determined the PCA index score. (1) Response bias was defined as the difference between lever presses and magazine entries, expressed as a proportion of the total responses [(lever presses−magazine entries)/(lever presses+magazine entries)]. (2) Latency score was calculated as the difference between the latency to approach the lever and the magazine after CS presentation; this difference was normalized by dividing by the maximum 8 s latency [(magazine latency−lever latency)/8]. (3) Probability difference was calculated as the difference between the probabilities of pressing the lever during the CS (i.e., the number of trials with a lever press out of 25 trials) minus the probability of entering the magazine. The PCA index score was the average of the response bias score, latency score, and probability difference. The values of this score ranged from 1.0 to −1.0, with a score of 1.0 indicating approaches and contacts of the lever on every trial, and a score of −1.0 indicating approaches and contacts of the magazine entry on every trial. Rats with an averaged PCA index score from PCA sessions 4 and 5 ranging from −1.0 to −0.4 were defined as GTs (i.e., rats more likely to direct behavior toward the food magazine than the lever), and rats with a PCA index score between +0.4 and +1.0 were designated as STs (i.e., rats more likely to direct behavior toward the lever-CS than the food magazine). Approach responses (response probability, number of contacts, and latency) were analyzed with repeated-measures ANOVAs with phenotype (STs, GTs) as the between-groups measure and training day as the within-subject factor. Rats that scored between -0.4 and +0.4 (‘Intermediates;’ INs) were not used in subsequent MCMCT experiments.

#### Surgery

Rats then underwent infusion surgeries, receiving bilateral microinjections of the AAV inhibitory virus (BF) or an inactive control plasmid, as well as neurotoxin 6-hydroxydoamine (6-OHDA) or sham infusions (striatum), followed by three weeks recovery, followed by a second surgery for implantations of optogenetic LED cannulae into the BF.

Rats were placed in vaporization chambers and anesthetized with 4–5% isoflurane delivered at 0.6 L/min O2 using a SurgiVet Isotec 4 Anesthesia Vaporizer until the animals were no longer responsive to a tail pinch and exhibited no hind limb withdrawal reflex. Heads were shaved using electric clippers and cleaned with a betadine scrub and alcohol pad. The animals were then mounted to a stereotaxic instrument (David Kopf Instruments) and isoflurane anesthesia was maintained at 1–3% for the remainder of the surgery. An ∼2.5 cm incision was made down the midline of the scalp to expose the skull. The animals’ body temperature was maintained at 37°C using Deltaphase Isothermal Pads (Braintree Scientific). Ophthalmic ointment was used for eye lubrication. To prevent hypovolemia and hemodynamic instability, 1 mL/100 g 0.9% NaCl was administered subcutaneously. Animals also received an injection of an analgesic (carprofen; 5.0 mg/kg, s.c; Henry Schein Medical) before surgery and once or twice daily as necessary for 48 h postoperatively for pain relief.

The optogenetic viral vector containing plasmid pAAV-nEF-Con/Foff 2.0-iC++-EYFP was obtained from AddGene (Watertown, MA; AddGene plasmid #137156-AAV8). The adeno-associated virus (AAV) coding for a NEF-driven, Cre-dependent expression of chloride-conducting channelrhodopsin iC++-EYFP was stereotaxically infused into the BF of 41 rats (N = 25, inhibitory AAV + 6-OHDA striatum lesions, and N = 16, inhibitory AAV + control sham striatum), targeting primarily the fronto-medial cortical projection systems arising from the HDB. A control (inactive) vector containing plasmid pAAV-Ef1a-Con/Foff 2.0-EYFP (AddGene plasmid #137162) was infused into the BF of 16 rats (control AAV + 6-OHDA striatum lesions).

One µl of pAAV-nEF-Con/Foff 2.0-iC++-EYFP or pAAV-Ef1a-Con/Foff 2.0-EYFP (control) vector was infused (bolus) into the BF at two sites per hemisphere (AP −0.6; ML ± 2.4; DV: −7.6; AP −1.0; ML ± 2.9; DV: −8.0 mm). The injector was left in place for 8 min to minimize diffusion into the injector tract. In the same surgery, rats received bilateral bolus infusions (4.0 mg/ml;1 ml/infusion) of the neurotoxin 6-hydoxydopamine (Sigma-Aldrich) or vehicle-only (1 mg/ml ascorbic acid in 0.9% NaCl) into two infusion sites (per hemisphere) of the dorsomedial striatum (AP +1.2; ML ± 2.2; DV: −4.5; AP +0.2; ML ± 2.5; DV: −5.0 mm). Rats were injected with desipramine hydrochloride (10 mg/kg, i.p.; Sigma-Aldrich) 30 min before 6-OHDA infusions for protection of noradrenergic neurons (Breese & Traylor, 1971). Rats were given 3 weeks to recover from the viral plasmid and 6-OHDA infusions.

#### Dual ACh/DA Disruption

In a second surgery, rats were anesthetized and prepared according to the parameters detailed above, and the LED cannulae with bilateral plastic fibers were implanted. Holes were drilled bilaterally on the skull approximately between the two viral infusion sites (AP -0.6 and AP -1.0) and the custom-sized cannulae were slowly lowered using a stereotaxic armpiece until the underside of the cannula units contacted the skull. Acrylic resin powder (Land Denture Mfg. Co. Inc.) and self-curing liquid resin (Ortho-Jet Liquid) were used to secure the cannulae in place as well as to adhere C-shaped 3D-printed plastic pieces for the attachment of clips to secure the receiver to the prongs of cannulae during traversals. Behavioral testing commenced after one week of recovery.

### LED Wireless Optogenetic System

In earlier pilot experiments on traditional optogenetic techniques using fiber optic cables to deliver laser illumination, it was found that connecting fiber optic cables to rats’ headstages during balance beam tasks could restrict their movement and disrupt performance, as well as being prone to lose connection if a rat accelerated quickly while traversing a rod, or if it fell from the MCMCT rods. Thus, a wireless LED optogenetic system was used for these experiments (Teleopto Wireless Optogenetic Stimulator System; Amuza Inc.; San Diego, CA). This light-delivering system comprised a wireless receiver (attached to the animals’ headstages), implanted custom-sized fiber optic LED cannulae (TelLCD-B-7.0-250-5.5, 470 nM, 5.59 mW; Amuza Inc.), a remote controller and an infrared emitter to initiate pulse generator LED stimulation (5 V) from a distance of up to 2 meters (STO mk II; Azuma Inc.). This wireless LED system has been used in previous studies to effectively modulate targeted neuronal activity and affect behavior (Chen et al., 2021; Dimitrov, 2024; Norman et al., 2021; Saito et al., 2025).

Rats were implanted with bilateral LED cannulae (470 nm) with plastic fibers, custom-sized to reach the basal forebrain region just above the optogenetic virus infusion sites (7.0 mm length; 5.5 mm between fibers; 250 µm fiber diameter). On test days, the wireless receiver (TeleR-1-P; 13 x 18 x 7 mm; weighing 1.4 g) was attached to connection prongs of the cannulae on the rats’ headstages and secured in place with custom-made clips. The wireless LED receiver was of minimal size and weight and when clipped to connection prongs did not cause weight distribution issues, impair balance, or restrict the movement of rats. The LED fibers were activated manually by a remote controller relay and pulse stimulator. Pulses were either 1s in duration or active for the duration of traversals in continuous stimulation runs.

#### MCMCT Test Battery

57 ChAT-Cre+ rats, 27 GTs (15 females) and 30 STs (20 females), were assigned pseudorandomly to three groups prior to surgeries: (1) inhibitory AAV + 6-OHDA (‘*Dual ACh/DA Disruption’*; N = 25; 7 GT females, 5 GT males; 8 ST females, 5 ST males); (2) inhibitory AAV + sham striatum (‘*ACh Disruption Only’*; N = 16; 5 GT females, 3 GT males; 5 ST females, 3 ST males); and (3) control AAV + 6-OHDA (‘*DA Lesion Only’*; N = 16; 4 GT females, 3 GT males; 6 ST females, 3 ST males). Following surgeries and recoveries, rats underwent a 17-day sequence of MCMCT sessions (see Table 2), beginning with a 3-day training block of runs on the plank, straight stationary (non-rotating) rod and straight rotating rod (Days 1-3). Next, rats traversed the straight rotating rod for a block of three consecutive sessions, receiving 1s inhibitory LED pulses (Days 4-6). Rats were then habituated to the zig-zag rod in two practice sessions, stationary on Day 7 and rotating on Day 8, followed by a block of three test days on the rotating zig-zag rod with 1s inhibitory LED pulses (Days 9-11). After a one-week break with no MCMCT sessions, rats underwent blocks of testing on the straight rotating rod (Days 12-14) and the zig-zag rotating rod (Days 15-17) as LED inhibition was delivered continuously throughout the duration of LED runs.

**Table 2.**
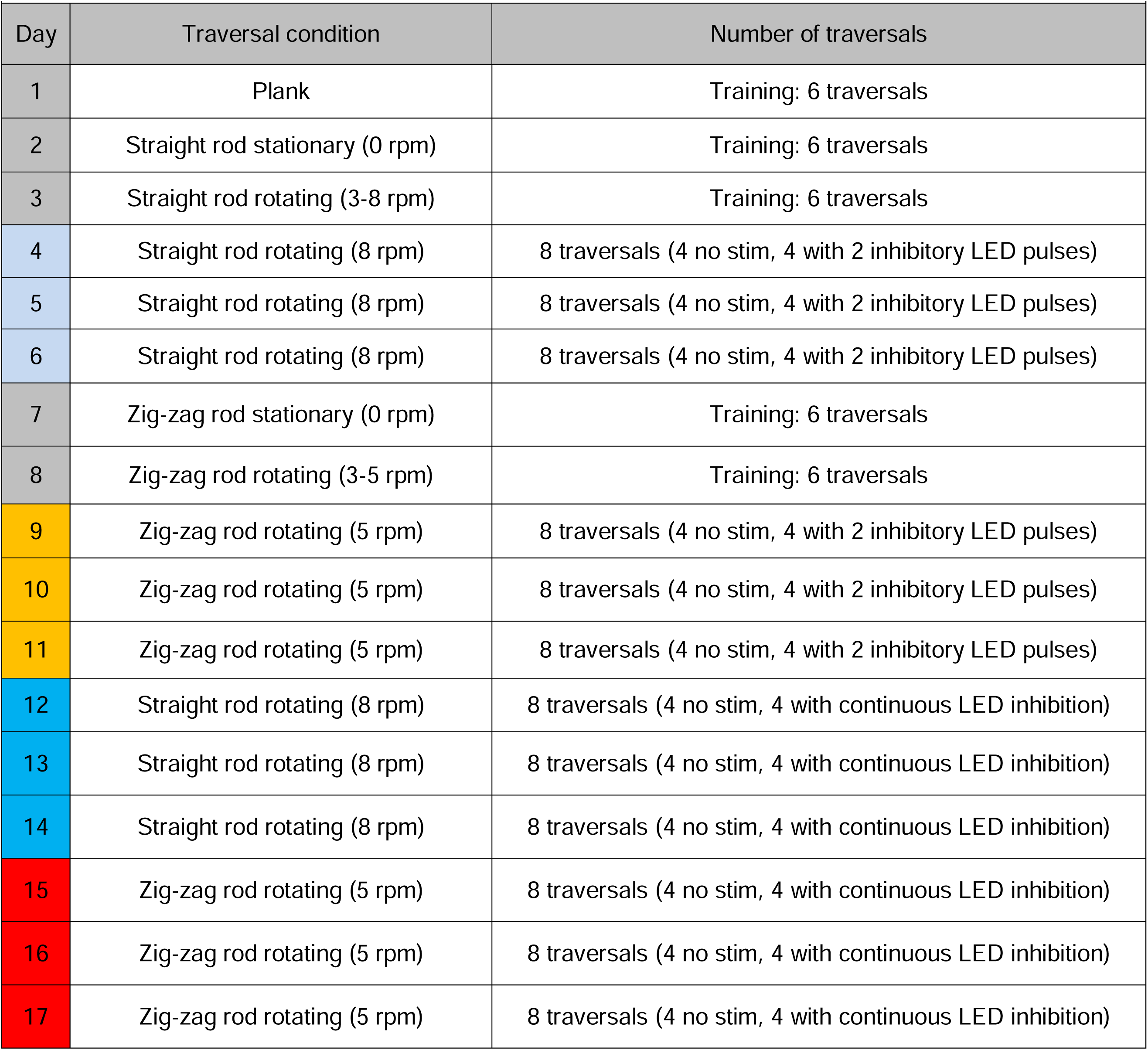
MCMCT Training and Test Sequence.

All rats that completed testing on the straight rotating rod with 1s LED pulses (Days 5-7) also completed testing on the straight rotating rod with continuous pulses (Days 9-11), except for 5 rats from early experiments that were only tested with 1s pulses on both the straight and zig-zag rods (all from group 1). Some rats were removed from zig-zag rod analyses due to complete inability or refusal to traverse. 1 *Dual ACh/DA Disruption* rat (male, ST) did not complete zig-zag runs with 1s LED pulses and 3 rats (1 *Dual ACh/DA Disruption* male GT, 1 *ACh Disruption Only* male GT and 1 *ACh Disruption Only* female *GT*) did not complete zig-zag runs with continuous LED pulses. In addition, rats with abnormally high baseline (no LED) fall rates on the zig-zag rotating rod (>0.75 falls per run) were removed from the analyses so that the effects of inhibitory LED pulses could be more clearly assessed. For 1s LED pulse runs, 9 such rats were removed: 2 *Dual ACh/DA Disruption* group rats (2 GT males), 5 *ACh Disruption Only* group rats (2 GT males, 2 ST males, 1 GT female), and 2 *DA Lesion Only* group rats (1 GT male, 1 GT female). For continuous LED runs, 17 such rats were removed: 8 *Dual ACh/DA Disruption* group rats (1 GT male, 2 ST male, 2 GT female, 3 ST female), 4 *ACh Disruption* Only group rats (1 GT males, 2 ST males, 1 GT female), and 5 *DA Lesion Only* group rats (1 GT male, 2 ST males, 2 ST females) (see Table 2). The final N values for each condition are provided in the Results section.

#### Optogenetic Testing Parameters

LED-induced optogenetic inhibition of BF cholinergic neurons was first assessed by delivering two 1s pulses per traversal on the straight rotating rod (Days 4-7). In this block of testing, rats performed 8 endbox-to-endbox traversals of the 3-meter rod per day, 4 with LED inhibitory pulses delivered twice per traversal, and 4 traversals in which no pulses were delivered. Pulses were activated manually by the experimenter when the rat crossed the 0.5-meter and the 1.5-meter demarcations enroute to the endboxes. Each pulse was 1s in duration. Thus, per test day, rats received 8 inhibitory pulses over 4 traversals (and for each 3 test-day block, 24 inhibitory pulses over 12 traversals). In the corresponding ‘no LED” traversals, no pulses were delivered. Pulses were given in two consecutive traversals – in traversals 1 and 2 and 5 and 6 (with no pulses in traversals 3 and 4 and 7 and 8) or traversals 3 and 4 and 7 and 8 (with no pulses in traversals 1 and 2 and 5 and 6). The order was randomized across all animals and conditions.

### Balance Beam task: the Michigan Complex Movement Control Task (MCMCT)

The MCMCT beam traversal apparatus (Kucinski et al., 2013; Kucinski & Sarter, 2021) was used to assess traversal performance measures in rats performing attention-demanding crossings of a ‘straight’ narrow rod or an angled rod surface (‘zig-zag’). The rods were 3.0 m in length (square; side lengths 1.59 cm), made from aluminum tubing covered with gray gaffer’s tape for traction. There was also a plank surface (13.3 cm width) placed directly on top of the straight rod and fitted firmly in place inside edges of the support towers used to habituate rats to traverse from endbox to endbox. The straight and zig-zag rods could be rotated using a 12 VDC electric motor controlled remotely by a pulse width modulator which was able to adjust the speed of rotation (up to 10 RPM) and switch the direction of rotation. A safety net was suspended 20 cm under the beam to catch the rats during falls. Two identical endbox stations were situated on top the support towers at opposite ends of the beam. These stations consisted of a 30 x 25 cm platform and were surrounded by a retractable wall structure (23 cm height in raised position) to allow conversion from an open platform to a boxed structure. On the wall facing the beam of each box structure were 9-cm wide openings that allowed the rats access to and from the beam. The walls were raised and lowered mechanically with a 12 VDC electric motor actuated remotely by a toggle switch. Lowering the walls and thereby turning the box into an open field platform was sufficient for rats to initiate beam traversal to enter the opposite endbox (with walls up). Between traversals, the walls of the endbox remained in the up position for 45 s. Videos of traversals were recorded with 8 bullet Ubiquiti Networks UVC-G4-PRO UniFi cameras (50P frame rate/8 MP resolution/3840×2160/3x zoom) mounted to the net frames parallel to both sides of the beam, 4 per side, to provide visualization of rats from both sides during traversals. Also attached to one side of the net frame, adjacent to the cameras, were LED indicator lights (red) activated by the pulse modulator to signal on video the precise timing of the pulse deliveries. The camera feeds were transmitted to a PC using a video capture system (Inmotion Studios, Ann Arbor, MI; UNVR with 8TB HDD Network Video Recorder Pro, 16 port giga ethernet POE switch, WD surveillance hard drive Raid1, cloud controller for UnFi camera system) and visualized in two continuous feeds (left and right sides) using customized video software (Arbor Computers, Ann Arbor, MI).

#### Straight rotating rod

Falls and traversal time were the measures of traversal performance on the straight rotating rod. These measures were analyzed in the same manner as previously described (Kucinski et al., 2019; Kucinski & Sarter, 2021). During falls, rats either fell into the net and were placed back onto the rod or, upon an imminent fall, the experimenter assisted the rat in regaining balance on the rod, and no further falls were counted until the rat regained a balanced posture and resumed forward movement. In addition to complete falls from the straight rod, instances in which the rats ceased forward movement and assumed a slouched, nearly immobile position on the rod were counted as falls. In either case, traversal time was corrected for fall-related disruptions of forward movement. In most instances, rats were able to regain forward movement within 2 s of being placed back onto the rod. Since the 1s LED pulses were not delivered until rats reached 0.5 m along the 3.0 m rod, falls that occurred prior to that point were not counted in both LED and no LED runs, and counting of traversal time commenced at the 0.5 m mark along the beam. In runs with continuous LED inhibition, performance measures were scored over the duration of the traversals. Also, since the 1s pulses were delivered twice per run (when rats approached 0.5 m and 1.5 m along the rod), a maximum of one fall per pulse or section of the rod (from 0.5 m to 1.5 m or 1.5 m to 3.0 m) was counted. Thus, a maximum of 2 falls per full traversal could be scored. To maintain consistency between testing conditions, a maximum of 1 fall per section of the rod (and 2 for each run) was also scored for continuous pulse runs. We previously reported high inter-rater reliability for extracting the behavioral measures from video recorded MCMCT runs (Avila et al., 2020).

#### Zig-zag rotating rod

A particularly challenging zig-zag rod variation was designed to determine the propensity for falls as rats performed traversals that taxed additional attentional control and flexibility of movement (Kucinski et al., 2018; Kucinski et al., 2019; Kucinski & Sarter, 2021). This rod featured two zig-zag sections, each 0.6 m in length, placed 0.6 m from either end of the rod and separated by a 0.6-m long straight rod in the middle. Each zig-zag section included three angled sections of varying lengths (two 10.7 cm and one 21.5 cm in length) that bended at 45° angles from the horizontal plane of the rod and connected to two straight sections (each 15 cm in length) that rested 3 cm above or below the plane the vertical plane of the rod. The diameter of the rod tube remained 2.54 cm² across the entire rod. The 1s LED pulses were delivered as the rats initiated forward movement onto the bent sections (2 per run). Falls and traversal time were the performance measures assessed on the zig-zag rotating rod runs. As done previously (Kucinski et al., 2019), one fall maximum was scored per zig-zag section (2 per run).

#### Scoring behavioral performance

Fall propensity from the straight rod was assessed by determining if a fall occurred during the period in which the rat first received an inhibitory pulse until either the rat received a second pulse (from the 0.5-meter to 1.5-meter demarcation) or until the end of each traversal (from the 1.5-meter demarcation to the endbox). The same criteria were used to count falls when no pulses were delivered. Thus, per traversal, there was a maximum of 2 falls scored, per test day a maximum of 8 falls, and per 3-day test block a maximum of 24 falls in LED runs. Fall totals over the 3-day block were scored out of a possible of 24 falls with inhibitory LED pulses and 24 possible falls with no pulses. In the next block of traversals, rats crossed the zig-zag rotating rod. Pulses were delivered when the front paws of the rats stepped onto the zig-zag sections as the rat moved toward the endbox, at approximately 1 and 2 meters along the rod. Fall totals were similarly scored with a maximum of 1 fall per pulse (or per zig-zag section), two per traversal, 8 per test day, and 24 for the block (with a maximum of 24 possible falls in no LED runs). Falls were also counted with continuous LED inhibition in two 3-day blocks, one on the straight rotating rod and one on the zig-zag rotation rod. For these runs, the inhibitory LED was activated for the entire duration of traversal - from when the rats first initiated traversal onto the rod until stepping into the endbox on the far end of the rod. Falls were counted in the same manner as the 1s pulse runs, with a maximum of two per traversal, one in first part of each run (either from the 0.5 to 1.5-meter demarcations on the straight rod or across the first zig-zag portion) and one in the second part (either from the 1.5 demarcation to the endbox on the straight rod or the across the second zig-zag portion). A maximum of 24 falls in LED inhibition and no LED runs could be scored for continuous inhibition runs as was in runs with 1s pulses.

### Histological visualization and quantification of AAV transfection space and 6-OHDA lesions

Following the end of all MCMCT tests, rats were deeply anesthetized with a lethal dose of sodium pentobarbital (270 mg/kg, i.p.) and transcardially perfused with phosphate buffer solution (PBS), followed by 4% paraformaldehyde in 0.15 M sodium-phosphate solution, pH 7.4. Brains were extracted and post-fixed in 4% paraformaldehyde for 24 hrs, rinsed with PBS and then placed in 30% sucrose solution until they sank. Brains were sectioned into 35-µm thick slices using a freezing microtome (CM 2000R; Leica) and stored in cryoprotectant until further histological processing.

To determine the expression space of the AAV in the BF, sections were double stained to both amplify the signal of the eYGFP fluorescent tag as well to visualize cholinergic neurons, using antibodies against the vesicular acetylcholine transporter (VAChT; Arvidsson et al., 1997). Sections were put in 0.1 M PBS (pH 7.3-7.4) overnight the day before staining. The next day sections were rinsed 3 times for 5 mins in 0.1 M PBS, then blocked in 0.1 Triton X100 diluted in PBS for 15 mins. Sections were then rinsed in PBS 3 times for 5 mins and incubated with 1% Normal Donkey Serum (NDS) and 1% Triton X100 made in PBS for 60 min at RT. The sections were then incubated overnight in the primary antibodies (chicken anti-GFP; 1:2000; Abcam, ab167453; and goat anti-VAChT; 1:500; Synaptic Systems, Cat. No. 139 103) in a cold room. The following day, sections were first rinsed 3 times for 5 each with PBS, and then incubated in secondary antibodies (Alexa 488-conjugated donkey anti chicken; 1:300; Jackson Laboratories, Cat. No. 703-545-155) and Alexa 594-conjugated donkey anti rabbit; 1:500; Jackson Laboratories, Cat. No. 711-585-152) in PBS, 0.5% normal donkey serum and 0.5% TritonX100 for 90 mins at RT. Sections were rinsed with PBS 3 times for 5 mins and then mounted and cover-slipped with Vectashield Antifade Mounting Medium (H-1000; Vector Laboratories). A Zeiss LM 700 confocal microscope was used to inspect and photomicrograph fluorescent neurons at 10X (for cell count estimates) and 20X (for verifying and documenting double-labeled cells) at two A-P levels (−0.72 and -0.96 mm). The microscope was equipped for sequential multi-track acquisition with 488 nm and 561 nm excitation lines and filter sets specific for Alexa 488 (Zeiss filter set 38 HE) and Alexa 594 (Zeiss filter set 54 HE), respectively. Single- and double-labeled neurons were counted in three subregions of the basal forebrain (nucleus basalis of Meynert, nbM; horizontal limb of the diagonal band, HDB; substantia innominata, SI). Using the ImageJ multipoint tool, superimposed counting frames were generated for each subsection (approximate sizes, actual subregion areas were based on irregular shaped outlines from the rat brain atlas: nbM: 0.4 mm x 0.8 mm; HDB:1.4 mm x 1 mm; SI: 0.5 mm x 0.9 mm) and counts were restricted to these frames. Counts were generated based on 2 sections per rat (−0.80 and -1.00 mm from bregma), yielding 4 counts per rat and subregion. Correlations of counts with falls were most apparent in the more posterior (−1.00 mm) sections, thus these are reported in the results section. Tile scanning/stitching was utilized at the 10X (4×4 tiles, 2368.77 by 2368.77 µm). The split view on Zen Black software was used to view single channels of the multi-track shots. Cells with green (Alexa 488) and red (Alexa 594) fluorescence located primarily in the soma were used in counts for double-labeled neurons.

To visualize the size and extent of 6-OHDA lesions, TH immunostaining was performed using a primary antibody (rabbit anti-TH; ab112, abcam), Vectastain Elite ABC Kit (PK-6101, Vector Laboratories) and DAB Substrate Kit (SK-4100, Vector Laboratories). Sections were first rinsed 3 times for 3 mins with 0.1 M PBS (pH 7.4) and then incubated in 0.3% H2O2 for 30 min. Sections were rinsed 3 times in 0.5% Triton (in PBS) for 3 mins each and then incubated in a blocking buffer (Vectastain buffered NGS + 0.3% Triton in PBS) for 1 hr at RT. Sections were then rinsed 3 times for 3 mins in PBS and incubated for 24-48 hrs with primary antibody (1:1000; 0.3% Triton in PBS) in a cold room. Sections were then rinsed 3 times for 3 mins with 0.3% Triton in PBS, incubated for 30 mins in secondary antibody (biotinylated goat-anti rabbit; 1:500, Vectastain) and then washed in PBS for 3 mins in 3 rinses. Sections were then incubated in ABC Elite (1:1000) for 30 mins. After another 3 rinses in PBS for 5 mins each, sections were developed in DAB substrate solution (buffer, DAB, H2O2, nickel) until they appeared dark (2-5 mins), and then rinsed 3 times in PBS for 5 mins. Sections were mounted with gelatin on SuperFrost Plus slides and allowed to dry overnight. The next day, the slides were dehydrated in an increasing alcohol series (70%, 95%, and 100%) and defatted in xylenes before coverslipping. Sections were inspected, and images were taken with a Leica DM400B digital microscope.

Photographs of TH immuno-stained sections in Group 1 rats (inhibitory AAV + 6-OHDA) taken at 1.25X from sections at approximately 500 µm posterior and anterior from 0.2 AP to 1.2 AP - the sites of infusion. The extent and location of the bleaching of TH stains was rated. TH-IR losses greater than 2.00 mm (dorsal-ventral plane) x 0.5 mm (medial-lateral plane) x 0.5 mm (anterior-posterior plane) were assigned the highest score (5). Lower scores (≤3) were assigned to smaller lesion sizes. We previously determined that TH losses in the mediodorsal part of the striatum, overlapping with the target field of prefrontal cortical projections are necessary and sufficient to produce falls in rats with BF cholinergic lesions (Kucinski et al., 2013; Kucinski et al., 2015). TH losses that were centered closest and that were mostly restricted to the mediodorsal caudate nucleus were assigned the highest score (5), whereas lesions spreading more laterally received lower scores. The size and placements scores for both hemispheres were averaged to a single DA lesion score (maximum of 5) for each rat.

### Statistical analyses

The effects of inhibitory 1s or continuous pulses on MCMCT performance measures were compared between the three disruption groups (inhibitory AAV + 6-OHDA or ‘*Dual ACh/DA Disruption’;* inhibitory AAV + sham striatum or ‘*ACh Disruption Only’*; control AAV + 6-OHDA or ‘*DA Lesion Only’*) using mixed measures ANOVA. For the primary comparisons of traversal performances between groups, Disruption Group was the between-subjects factor and stim (LED or no LED) was the within-subjects factor. Between-subjects factors sex and Phenotype (ST or GT) and within-subjects factor Day (within each 3-day block) were used when applicable. Tukey’s Honest Significant Difference test was used for post hoc comparisons following significant main effects or interactions. Pearson’s correlation coefficient (r value) is provided for linear correlations between histological quantifications and performance measures. Statistical analyses were performed using SPSS for Windows (version 25: IBM SPSS). Assumptions underlying the statistical model were assessed. In cases of violation of the sphericity assumption, Huyhn–Feldt-corrected *F* values, along with uncorrected degrees of freedom, are given. Alpha was set at 0.05 except for the analysis of falls. Exact *P* values are reported as recommended previously (Greenwald et al., 1996).

## Results

### Straight Rotating Rod, 1s LED inhibitory pulses

The effects of optogenetic BF cholinergic inhibition on MCMCT performance measures were first assessed over a 3-day block with the straight rotating rod in 57 rats (*Dual ACh/DA Disruptio*n group: N = 25, *ACh Disruption Only* group: N = 16, *DA Lesion Only* group: N = 16). 1s LED inhibitory pulses were delivered twice per traversal in stim runs, and no pulses were given in baseline ‘no LED’ runs. In this and subsequent analyses, GT/ST status and F/M sex were not significant factors or interactions, and so for simplification F/M rats and GT/ST rats were combined, and only the factors stim (LED or no LED inhibition) and Disruption Group are reported. Since a maximum of 1 fall was scored per 1s pulse (or per section of the rod), there were a maximum of 8 falls per day over the 3-day block, and thus total falls per block are reported out of a maximum of 24. Figure 1A-D shows the number falls by each individual rat within each traversal task condition. Traversal times are reported per run (one endbox to endbox traversal).

**Figure 1.**
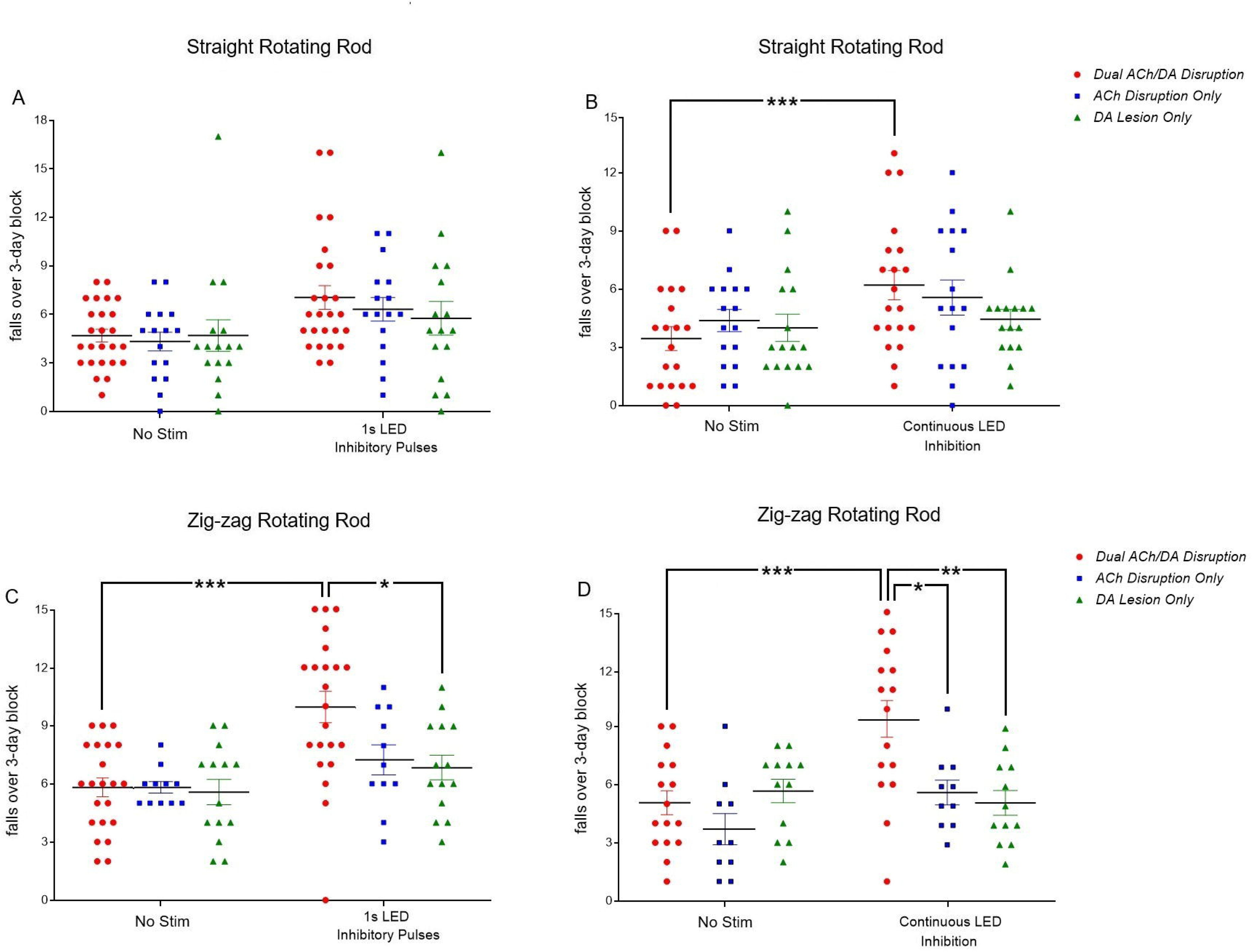
Falls by Disruption Group with no LED or LED inhibition. The combined 3-day total out of a possible 24 falls in each test block is shown per rat. (A) Straight rotating rod, 1s LED inhibitory pulses (N = 57). Falls were elevated overall in LED runs but there were no group differences. (B) Straight rotating rod, continuous pulses (N = 52). Falls were again elevated with continuous LED inhibition, and falls were increased relative to no LED trials only in the *Dual ACh/DA Disruption* group; pairwise post hoc comparison following significant LED stim x group interaction. (C) Zig-zag rotating rod, 1s LED inhibitory pulses (N = 47) and (D) Zig-zag rotating rod, continuous LED inhibition (N = 37). Falls by rats with *Dual ACh/DA Disruption* were elevated with 1s LED inhibitory pulses compared to *DA Lesion Only* rats, and elevated with continuous LED inhibition relative to *ACh Disruption Only* and *DA Lesion Only* group rats. *P<0.05, **P<0.01, ***P<0.001; post hoc effect of LED stim after significant Disruption Group x LED stim interaction.

Falls were overall significantly elevated from the straight rotating rod by optogenetic inhibitory 1s LED pulses (Figure 1A; effect of stim: F(1,54) = 32.65, p < 0.001; no LED: 4.58±0.36 falls per rat over the 3-day block, LED: 6.47±0.48), although this effect of stim did not interact significantly with Disruption Group (F(2,54) = 1.53, p = 0.23; group means: *Dual Loss ACh/DA:* no LED 4.68±0.39 falls, LED: 7.04±0.73; *ACh Loss Only* group: no LED: 4.31±0.58, LED: 6.31±0.72; *DA Lesion Only* group: no LED: 4.69±0.97, LED: 5.75±1.04. Falls in general progressively declined over the 3-day block of 8 traversals per day (effect of Day: F(2,108) = 21.45, p < 0.001; falls on Day 1 (4.88±0.44) were higher than on Day 2 (3.40±0.28; p < 0.001) and Day 3 (p < 0.001), and falls on Day 2 were higher than Day 3 (p = 0.01); interactions involving Day and Disruption Group were non-significant, F<1.87, p>0.12 for all). In subsequent conditions, falls generally declined in later test days within each block, however since there were no significant interactions involving Day and Disruption Group, these are not reported.

Traversal time was slower in runs with LED pulses (effect of stim: F(1,54) = 29.88, p < 0.001; no LED: 11.71±0.54 s per run; LED: 12.89±0.64), however there was not a significant interaction between LED stim and Disruption Group (F(2,54) = 0.14, p = 0.94; group means: *Dual ACh/DA Loss*: no LED: 11.74±0.65 seconds per traversal, LED: 12.91±0.89; *ACh Disruption Only:* no LED: 11.08±1.15 s, LED: 12.29±1.33; *DA Lesion Only:* no LED: 12.30±1.23 s, LED: 13.44±1.28).

### Straight rotating rod, continuous LED inhibition

Traversal performances were then assessed with continuous LED inhibition over the duration of traversals in stim runs (Figure 1B; N = 52 rats; *Dual ACh/DA Disruptio*n group: N = 20; *ACh Disruption Only*: N = 16; *DA Lesion Only*: N = 16). As with 1s LED pulses, continuous LED inhibition overall elevated falls (effect of stim: F(1,49) = 23.09, p < 0.001; no LED: 3.90±0.36 falls, LED: 5.46±0.44) and this effect of stim interacted significantly with Disruption Group (F(2,49) = 5.35, p = 0.008); however, post hoc analysis indicated that falls did not significantly differ between groups in either no LED or LED stim trials (no LED: F(2,51) = 0.56, p = 0.57; LED stim: F(2,51) =1.43, p = 0.25; group means: *Dual ACh/DA Disruption:* no LED: 3.45±0.61 falls, LED: 6.20±0.76; *ACh Disruption Only:* no LED: 4.38±0.57, LED: 5.56±0.90; *DA Lesion Only*: 4.00±0.70, LED: 4.44±0.52). Although there was not a significant Disruption Group effect in LED stim trials, within the *Dual ACh/DA Disruptio*n group pairwise comparisons between no LED and LED stim trials indicated a significant increase in falls with LED stim (F(1,19) = 57.47, p < 0.001), while the marginal increase in falls by the *ACh Disruption Only* group with LED stim was not significant (F(1,15) = 3.70, p = 0.07), and LED inhibition failed to have any effect in the *DA Lesion Only* group (F(1,15) = 0.73).

Traversal time overall was significantly slowed in LED inhibition trials (effect of stim: F(1,49) = 4.91, p = 0.03; no LED: 8.92±0.51 s per run, LED: 9.49±0.63), however the stim x Disruption Group interaction was non-significant (F(2,49) = 1.34, p = 0.27; group means: *Dual ACh/DA Disruption* group: no LED: 8.71±0.82 s, LED: 9.64±1.09; *ACh Disruption Only* group: no LED: 8.28±0.81, LED: 8.27±0.79; *DA Lesion Only* group: no LED: 9.84±1.05, LED: 10.53±1.29).

### Zig-zag rotating rod, 1s LED inhibitory pulses

Falls and traversal times were next assessed using the more challenging zig-zag rotating rod (N = 47 rats; *Dual ACh/DA Disruptio*n group: N = 22, *ACh Disruption Only* group: N = 11, *DA Lesion Only* group: N = 14). Presenting 1s Ach inhibitory LED pulses increased falls overall (effect of stim: F(1,44) = 26.48, p < 0.001; no LED: 5.74±0.30 falls, LED: 8.40±0.50), and this effect interacted significantly with Disruption Group (F(2,44) = 5.26, p = 0.009). Falls significantly differed between groups in LED stim trials (F(2,46) = 5.01, p = 0.01) but not in the trials with no LED (F(2,46) = 0.07, p = 0.94). Tukey HSD post hocs indicated that *Dual ACh/DA Disruption* group rats fell more in 1s LED pulse trials than *DA Lesion Only* group rats (p = 0.02) but the higher falls vs the *ACh Disruption Only* did not reach significance (p = 0.07) (Figure 1C; group means: *Dual ACh/DA Disruption*: no LED: 5.82±0.48 falls, LED: 9.95±0.80; *ACh Disruption Only* group: no LED: 5.82±0.30, LED: 7.27±0.78; *DA Lesion Only* group: 5.57±0.65, LED: 6.86±0.50). Pairwise post hoc comparisons within each group showed a significant increase in falls with LED pulses in *ACh/DA Disruption* rats (F(1,21) = 34.58, p < 0.001) and no such effect in *ACh Disruption Only* (F(1,10) = 3.49, p = 0.09) or *DA Lesion Only* rats (F(1,13) = 3.63, P = 0.08).

Overall, rats’ traversal times were slowed by LED inhibition (effect of stim: F(1,44) = 9.54, p = 0.003), however the factors of stim and Disruption Group did not interact significantly (F(2,44) = 1.73, p = 0.19; means: *Dual ACh/DA Loss* group: no LED: 20.25±1.49 seconds per traversal, LED: 21.08±1.58; *ACh Disruption Only* group: no LED: 18.61±2.13, LED: 19.30±2.44; *DA Lesion Only* group: no LED: 27.65±2.67, LED: 30.11±2.84).

### Zig-zag rotating rod, continuous LED inhibition

With continuous LED inhibition in stim runs on the zig-zag rotating rod (Figure 1D; N = 37 rats; *Dual ACh/DA Disruption*: N = 16, *ACh Disruption Only*: N = 10, *DA Lesion Only*: N = 11), falls were significantly increased in LED trials compared to no LED trials (effect of stim: F(1,34) = 12.35, p = 0.001; no LED: 4.95±0.40 falls, LED: 6.82±0.63) and this effect varied between groups (stim x Disruption Group interaction: F(2,34) = 9.55, p = 0.001). As with 1s LED inhibitory pulses on the zig-zag rotating rod, the groups differed significantly in trials with LED inhibition but not in trials without LED inhibition (no LED: F(2,36) = 2.32, p = 0.13; LED inhibition: F(2,46) = 7.87, p = 0.002). Tukey HSD post hocs showed more falls in the *Dual ACh/DA Disruption* group than both the *ACh Disruption Only* (p = 0.02) and *DA Lesion Only* (p = 0.003) groups with LED inhibition (group means: *Dual ACh/DA Disruption*: no LED: 5.06±0.62 falls, LED: 9.44±1.00: *ACh Disruption Only*: no LED: 3.70±0.80, LED: 5.70±0.63; *DA Lesion Only*: no LED: 5.91±0.61, LED: 5.00±0.73). Within groups pairwise comparisons indicated significantly higher falls with LED inhibition in *Dual ACh/DA Disruption* rats (F(1,15) = 23.00, p < 0.001) but higher falls in *ACh Disruption Only rats* did not reach significance (F(1,9) = 4.87, p = 0.06) and there was no detectable effect of LED inhibition in *DA Lesion Only* rats (F(1,10) = 1.79, p = 0.21).

Overall, traversal times were slower in stim LED inhibition runs (effect of Stim: F(1,34) = 5.10, p = 0.03), however the Disruption group x stim interaction was non-significant (F(2,34) = 0.34, p = 0.72; group means: *Dual ACh/DA Disruption*: no LED: 17.97±2.06 s, LED: 19.56±2.23; *ACh Disruption Only*: no LED: 16.17±3.28, LED: 17.38±3.42; *DA Lesion Only*: no LED: 22.00±3.40, LED: 22.62±3.42).

### Lack of Sign Tracker vs Goal Tracker differences

Potential GT/ST differences on falls and traversal times were assessed in a separate analysis, using stim (LED inhibition or no LED) as the within-subjects factor, and GT/ST, sex, and Disruption Group (*Dual ACh/DA Disruption*, *ACh Disruption Only,* or *DA Lesion Onl*y) as between-subject factors. GT and ST rats did not differ in falls on any rod condition, nor were there any significant 2,3 or 4-way interactions involving GT/ST as a factor with stim or sex (F<2.33, p>0.11 for all). In the *Dual ACh/DA Disruption* group, falls did not differ between GTs and STs from the straight rod with 1s LED inhibitory pulses (effect of GT/ST: F(1,23) = 0.02, p = 0.91) nor on the straight rod with continuous LED inhibition (effect of GT/ST: F(1,18) = 0.75, p = 0.13), and the stim x GT/ST interaction was non-significant in both conditions (F<2.50, P>0.13 for both). Similarly, on the zig-zag rotating rod, falls did not differ between GTs and STs or interact with stim with either 1s LED inhibition (GT/ST effect: F(1,20) = 0.01, p = 0.98; stim x GT/ST: F(1,20) = 0.47, p = 0.50) nor with continuous LED inhibition (GT/ST effect: F(1,14) = 0.16, p = 0.70; stim x GT/ST: F(1,14) = 0.16, p = 0.69). Figure 2A shows the number of falls from the zig-zag rotating rods for all GT and ST rats in the *Dual ACh/DA Disruption* group. 1s LED inhibition increased falls by GTs (F(1,9) = 10.96, p = 0.009; no LED: 6.10±0.69 falls per 3-day block, LED: 9.70±1.36 falls) and STs (F(1,11) = 23.71, p < 0.001; no LED: 5.58±0.69 falls, LED: 10.17±0.99 falls), and continuous LED inhibition also increased falls by both GTs (F(1,7) = 10.18, p = 0.02; no LED: 5.00±0.85 falls, LED: 9.00±1.34) and STs (F(1,7) = 11.54, p = 0.01; no LED: 5.13±0.95 falls, LED: 9.88±1.57).

**Figure 2.**
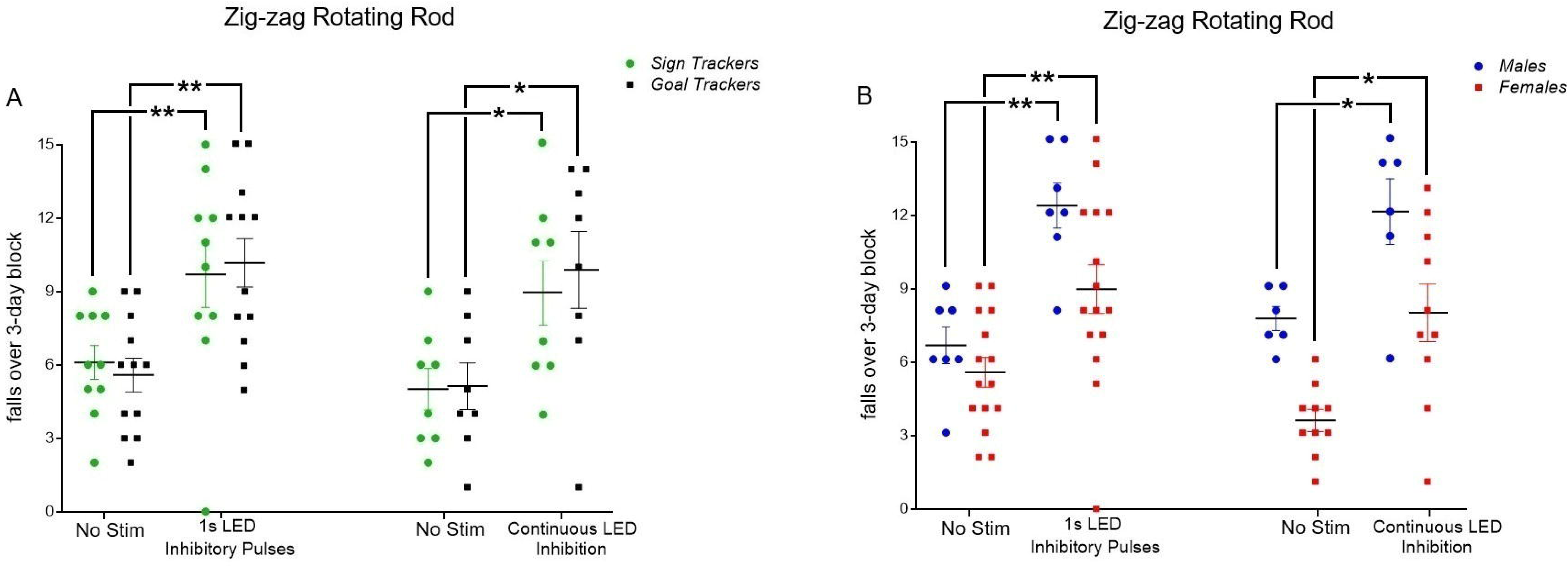
(A) Falls by GTs and STs in the *Dual ACh/DA Disruption* group from the zig-zag rotating rod with 1s LED inhibitory pulses (N = 12 GT, 10 ST) or continuous LED inhibition (N = 8 GT, 8 ST). Falls were elevated in both GTs and STs by both 1s LED pulses and continuous LED inhibition of BF cholinergic neurons. (B) Similarly, falls by both female and male rats in the *Dual ACh/DA Disruption* group were increased by both 1s LED pulses (N = 15 females, 7 males) and continuous LED inhibition (N = 10 females, 6 males) of BF cholinergic neurons. Male rats generally fell more than females across all conditions; however, this effect did not interact with Disruption Group. *P<0.05, **P<0.01 effect of LED stim within GT/ST phenotype or sex.

Traversal times also did not differ between GT and ST on any rod condition, nor where there any significant 2,3 or 4-way interactions involving GT/ST status with stim or with sex (F<2.46, p>0.10 for all).

### Sex Difference Effects

On the straight rod, females and males did not differ in falls, nor where there any significant interactions between sex and 1s LED or continuous LED inhibition (1s LED inhibition runs: effect of sex: F(1,51) = 1.93, p = 0.17; sex x stim, sex x Disruption Group, and 3-way interaction, F < 2.35, p > 0.13 for all; overall mean (Stim and no stim runs): females: 10.23±0.83 falls per 3-day block, males: 12.36±1.52 falls; continuous LED runs: effect of sex: F(1,46) = 1.48, p = 0.23; sex x stim, sex x Disruption Group, and 3-way interaction; F < 5.16, p > 0.11 for all; overall mean (stim and no stim runs): females: 8.50±0.93 falls per 3-day block, males: 10.55±1.16 falls).

However, on the more challenging zig-zag rod, larger males generally fell more often than females regardless of LED or no LED condition (1s LED inhibition runs: effect of sex: F(1,41) = 6.23, p = 0.02; sex x stim, sex x Disruption Group, and 3-way interaction, F < 1.69, p > 0.20 for all; overall mean (stim and no stim runs): females: 12.94±0.81 falls per 3-day block, males: 16.73±1.00 falls); continuous LED runs: F(1,31) = 15.87, p < 0.001; sex x stim, sex x Disruption Group, or 3 way; F < 1.37, p > 0.26 for all; overall mean (stim and no stim runs): females: 10.04±0.72 falls per 3-day block, males: 16.82±1.41 falls). Although males generally fell more than females from the zig-zag rotating rods, falls by both sexes in the *Dual ACh/DA Disruption* group were negatively impacted by 1s LED inhibitory pulses (7 males and 15 females) or continuous LED inhibition (6 males and 10 females) (Figure 2B; effect of stim P < 0.01 for all). The larger body size of male rats (417.53±32.42 g in 22 male rats vs 320.98±19.72 g in 35 female rats) was likely the primary cause of greater overall propensity of males to fall, as male rats have more trouble regaining upright posture following slips from the rotating rods.

The males generally performed slower traversals than females across the straight and zig-zag rotating rod conditions, and this was consistent across the Disruption Groups (effect of sex, F > 16.92, P < 0.001 for all; overall means (stim and no stim runs): straight rod: 1s LED inhibition: females: 10.85±0.72 s, males: 14.61±0.77 s; continuous LED inhibition: females: 7.83±0.67 s, males: 10.42±0.68 s; zig-zag rod: 1s LED inhibition: females: 19.29±1.35 s, males: 27.99±2.05 s; continuous LED inhibition: females: 15.39±1.70 s, males: 26.46±2.38 s; sex x Disruption Group interactions: F<2.13, p>0.13 for all).

### Histological Analysis

#### Basal Forebrain

Optogenetic AAV infusions targeting cortically projecting neurons of the basal forebrain were conducted using the same parameters as prior AAV infusion experiments using DREADDs (Kucinski et al 2019, Kucinski et al 2022). The inhibitory AAV spread diffusely into the basal forebrain: anteriorly to cholinergic and non-cholinergic neurons of the globus pallidus and into the ventral pallidum and bed nucleus of the stria terminalis; and posteriorly through the ventromedial globus pallidus and magnocellular cholinergic neurons of the nucleus basalis of Meynert (nbM) (Zaborszky et al., 2008), as well as the cholinergic neurons of the substantia innominata (SI) and the horizontal limb of the diagonal band (HDB). The LED fibers were placed into the basal forebrain at approximately AP -0.8, halfway between two infusion sites of the AAV (AP -0.6 and -1.0) (See Figure 3E for a schematic of the BF target area).

**Figure 3.**
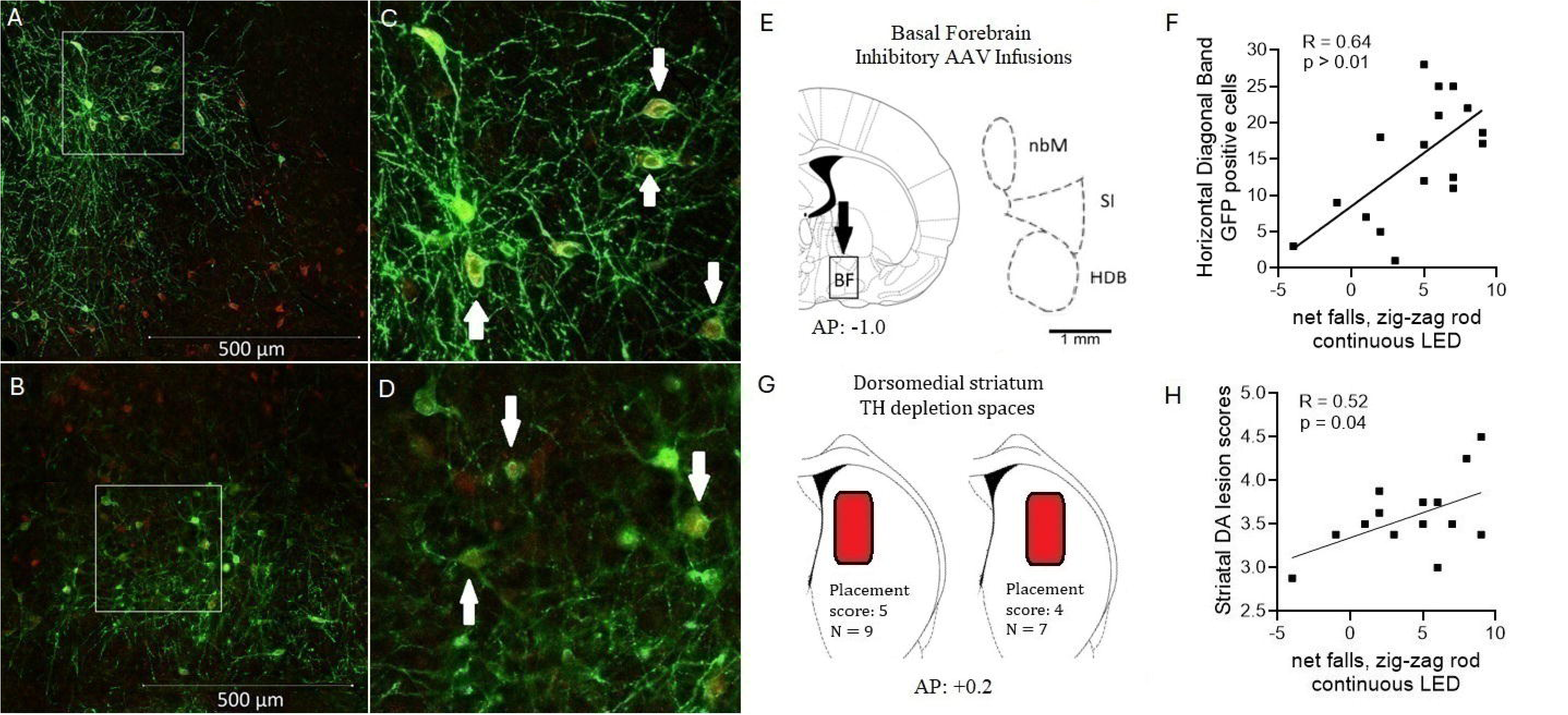
Visualization of the chloride-conducting channelrhodopsin iC++-EYFP in the BF (SI) of cholinergic and noncholinergic neurons (Alexa 594, appearing green, indicates the expression of the optogenetic viral construct (GFP amplified tag); Alexa 488, appearing red, indicates the presence of the VAChT in cholinergic cells). Low-magnification photomicrographs (500-μm scale inserted) of a coronal section showing the SI of a high falling rat with the inhibitory AAV + 6-OHDA lesion (15 falls in 24 trials on the zig-zag rod with continuous LED inhibition) (A) and a low falling rat with the inhibitory AAV + 6-OHDA lesion (B) (6 falls in 24 trials). These photomicrographs exemplify the dense presence of magnocellular cholinergic neurons (red) as well as numerous neurons expressing GFP or co-labelled cells. The areas marked by a whitish overlay in (A) and (B) are magnified in (C) and (D), respectively, and show several double-labeled neurons (arrows). More robust expression of the inhibitory AAV and stronger co-localization with cholinergic cells is seen with the high falling rat (A and C). A schematic of the inhibitory AAV infusion target in the BF at -1.0 AP is shown in (E). Boundaries of the cell counting areas for the 3 BF subregions (nBM, SI, HDB) is also shown. For the 16 *Dual ACh/DA Disruption* group rats included in the histologic analysis, viral transfection efficacy and the degree of double labeling was significantly correlated with counts in the SI as well as the HDB and nBM. In addition, the efficacy of the optogenetic virus activation, in terms of fall propensity (LED-induced falls minus no LED falls, zig-zag rotating rod), was significantly correlated with viral transfection efficacy in the SI and HDB (F), but not for counts obtained from the nBM. (G) shows a schematic of the dorsomedial striatum 6-OHDA infusion sites for the 16 *Dual ACh/DA Disruption* group rats at AP +0.2. The red shaded areas depict the TH depletion spaces directly in the dorsomedial stratum (9 rats) with placement scores of 5 (9 rats) and placement scores of 4 (7 rats) with depletion spaces slightly lateral. The combined TH rating scores (average of placement and size; size scores varied between 1 and 4; AP +1.2 and AP +0.2) correlated negatively with net falls (LED – no LED) from the zig-zag rotating rod with continuous LED inhibition (H).

Figure 3A-D shows microphotographs from the SI of two *Dual ACh/DA Disruption* group rats, one ‘high faller’ (15 falls in 24 trials from the zig-zag rod trials with continuous LED inhibition) and one ‘low faller’ (6 falls in 24 trials), exemplifying the widespread expression of the inhibitory AAV (Alexa 488; GFP, green). The magnified area shows cholinergic cells (Alexa 594; VAChAT, red) co-expressing the inhibitory AAV. The high faller had more robust expression of the inhibitory optogenetic AAV and stronger co-localization with cholinergic cells. Transfection efficacy of the AVV was estimated using manual counts of 1) GFP-positive cells and 2) co-labeled VAChAT-positive cells in three major regions of the BF projecting to cortical regions (nbM, SI, HDB) using the *Dual ACh/DA Disruption* group (N = 16) rats. Also, off-target expression of the AAV was assessed by determining the percentage of non VAChAT labeled cells with GFP expression in the BF subregions.

#### Neostriatal 6-OHDA lesions

Dopamine terminals within the dorsomedial striatal projection region of the medial prefrontal (prelimbic) cortex were targeted with 6-OHDA infusions using the same parameters as previous experiments (Avila et al., 2020; Kucinski et al., 2013; Kucinski et al., 2017; Kucinski et al., 2019). TH lesion scores were graded to characterize the size and placement of the TH depletion spaces (see Figure 3G for a schematic of target spaces). TH lesions sized >2.00 mm (dorsoventral) × 0.5 mm (mediolateral) × 0.5 mm (anteroposterior) were rated highest (5) and lesions smaller than 0.1 mm x 0.1 mm received the lowest score (1). Lesions centered in and largely limited to the dorsomedial region received the highest score (5), whereas lesions lateral to this prefrontal projection region received lower grades (see Methods for further details and Figure 3G). Placement scores of the lesions were consistently high, with an average of 4.64±0.11 (out of 5). Size scores were 2.55±0.19 (out of 5), and the average overall TH score was 3.59±0.10. The TH scores of 16 *Dual ACh/DA Disruption* rats significantly correlated with LED-induced falls from the zig-zag rotating rod with continuous LED inhibition (r = 0.52, p = 0.04), as in, rats with striatal TH depletion areas that tended to be more localized to the dorsomedial striatum and larger were more likely to fall with continuous LED inhibition (Figure 3H). TH scores did not correlate significantly with LED-induced falls from the straight rod with either 1s or continuous LED inhibition, nor from the zig-zag rod with 1s LED inhibition (r<0.26, p>0.33 for all).

### Relation of Histological Virus Analysis to Falls

Basal forebrain GFP cell counts generally correlated positively with net falls (falls in LED stim trials minus falls in no LED trials over the 3-day block) but were primarily revealed in performance on the zig-zag rod. In trials with the straight rotating rod, with both 1s LED inhibition and continuous LED inhibition correlations between GFP-positive or co-labeled GFP+VAChAT cell counts and net falls did not reach statistical significance (GFP-positive: 1s LED pulses: r < 0.50, p > 0.05 for all subregions; continuous LED inhibition: r < 0.45, p > 0.08 for all; co-labeled GFP+VAChAT: 1s LED pulses: r <0.27, p > 0.32 for all subregions; continuous LED inhibition: r < 0.30, p > 0.25 for all).

In trials with the straight rotating rod, with 1s LED inhibition correlations between GFP-positive or co-labeled GFP+VAChAT cell counts and net falls did not reach statistical significance (GFP-positive: 1s LED pulses: r < 0.50, p > 0.06 for all subregions; co-labeled GFP+VAChAT: 1s LED pulses: r <0.21, p > 0.08 for all subregions). With continuous LED inhibition, co-labeled GFP+VAChAT cells in the HBD only correlated with net falls (r = 34, p = 0.02; nBM and SI: r < 0.11, p > 0.22).

However, on the zig-zag rotating rod runs, LED inhibition with 1s pulses induced falls that correlated with GFP-positive cell counts in the SI (r = 0.56, p = 0.02; r<0.36, p>0.17 for the HDB and nbM), but not with the co-labeled cell counts in any subregion (r<0.20, p>0.08 for all). With continuous LED inhibition on zig-zag rod trials, GFP-positive cells correlated with net falls in the SI (r = 0.58, p = 0.02) and the HDB (r = 0.64, p = 0.01; Figure 3F) (nbM: r = 0.33, p = 0.21) and co-labeled cells correlated with net falls in the HDB (r = 0.59, p = 0.02; SI and nbM: r<0.51, p>0.05).

To assess off-target expression of the inhibitory optogenetic AVV, the percentage of GFP positive cells that were also VAChAT positive (co-labeled) were determined in each BF subregion: nBM: 89.77±1.02%, SI: 79.69±3.93% and HDB: 70.44±6.86% (79.96±2.85% total). This off-target expression may arise from leaky expression of the optogenetic AAV or be due to staining/visualization discrepancies. The HDB and SI contain 90-95% non-cholinergic neurons (Gritti et al., 2006), thus more non-targeted expression in those regions would be expected, however these values are low relative to their proportions of non-cholinergic cells in the BF. Also, in those regions were the strongest correlations between co-labelled cells and net falls by dual disruption group rats, thus the effects of potential off-target expression appear to be minimal but could conceivably have affected the results. Nonspecific DREADD inhibition of BF neurons was effective at negatively impacting falls and attentional processing in previous studies (Kucinski et al., 2019; Kucinski et al., 2022).

## Discussion

Cholinergic deficits in the cortex and BF are commonly observed in PD, particularly in individuals with high fall propensity, as well as cognitive dysfunction (Grothe et al., 2021; Schulz et al., 2018). Previous studies indicated that 6-OHDA lesions of dorsomedial striatal DA terminals combined with lesion-induced losses of cholinergic BF neurons, or nonspecific DREADD inhibition of BF neurons induced heightened falls in rats from the MCMCT rotating beams (Kucinski et al., 2013; Kucinski et al., 2017; Kucinski et al., 2019). Here, the dual DA and ACh disruption model was extended by combining optogenetic inhibition of ACh neurons in the BF with striatal 6-OHDA lesions to assess the effects of selective, phasic inhibitions of BF cholinergic neurons with dorsomedial striatal DA depletion. The more challenging zig-zag rotating rod which requires superior attentional control to effectively navigate (Kucinski et al., 2018) was added to better reveal balance and coordination deficits uncovered by the dual disruption model. These results confirm that selective optogenetic inhibition of cholinergic BF neurons, a manipulation that impairs sustained attention and disrupts cue detection and integration (Gritton et al., 2016), when combined with striatal DA depletion, induces a higher propensity to fall than either disruption alone, and this propensity is revealed especially on the more challenging zig-zag rod task.

Unlike rats with combined ACh and DA system disruption, rats with only ACh disruption and sham striatal infusions did not exhibit significantly elevated falls with BF cholinergic optogenetic inhibition, consistent with previous findings using lesions (Kucinski et al., 2013), validating the critical importance of dual striatal DA and cortical ACh systems in modulating complex movement. It was suggested that PD cognitive and related symptoms are a dual syndrome of DA mediated fronto-striatal disruption and a dementia syndrome in part mediated by cholinergic systems (Kehagia et al., 2013). There may be a heightened reliance on cholinergic-cognitive resources to guide movement to compensate for underlying deficits of striatal DA mechanisms in PD or after striatal DA lesions in rats. When the BF cholinergic system is also disrupted in such individuals, these cognitive resources can no longer be recruited effectively to compensate for striatal DA losses, unmasking deficits in controlling movement and resulting in falls. This diminished striatal control of complex movement may be particularly problematic in situations that demand focused attention and cognitive flexibility such as navigating complex surfaces or while dual-tasking (Kucinski et al., 2013; Sarter et al., 2014). The unmasking of underlying DA deficits in PD patients that also develop cholinergic losses may be a primary cause of falls and explain why L-DOPA therapy that enhances dopaminergic output does not improve falls. Pro cholinergic treatments in rat models (Kucinski et al., 2015; Kucinski et al., 2017) and human subjects (Albin et al., 2021; Quik et al., 2019; Stuart et al., 2020) suggest that such drugs may be a beneficial adjunct therapy against falls and related gait, balance, and postural impairments in PD patients with cognitive/cholinergic deficits.

Falls by rats with dual ACh and DA disruption became more negatively impacted in each successive block of their MCMCT tests with optogenetic stimulation parameters (see Figure 1A-D). Falls by the dual disruption group were not increased on the straight rotating rod by 1s LED inhibitory pulses, but were elevated by continuous LED inhibition on the straight rotating rod. On the more demanding zig-zag rotating rod, falls with 1s LED inhibitory pulses by the dual disruption group were more frequent than such falls by rats with DA lesions only or by the dual disruption group themselves on control non-LED trials. Continuous LED inhibition caused the dual disruption group to fall more than both the ACh inhibition-only group and the DA lesion-only group. Thus, falls were revealed by the dual system disruptions especially on the more challenging zig-zag traversal beam, and continuous LED inhibition appeared to have a slightly stronger and more consistent induction of falls than 1s LED pulses even on the less challenging straight beam. Studies have verified the greater effectiveness of longer trains (5-30s) at stimulation of BF cholinergic neurons using c-Fos expression and microdialysis (Zant et al., 2016) and *in vivo* electrophysiological recordings (Ma & Luo, 2012).

Marginal increases in falls by rats with the inhibitory optogenetic AAV and sham striatal lesions in LED trials compared to non-LED trials by the same rats approached but did not reach significance in 3 of the 4 conditions. In previous lesion studies, similar marginal increases in falls were induced in rats with only BF cholinergic lesions, specifically on more demanding traversal tasks, but falls by rats with dual BF cholinergic lesions together with the striatal DA lesions were robustly more frequent (Kucinski et al., 2013), just as here. Falls on the MCMCT were also induced by BF nonspecific neuronal inhibition using DREADDs (Kucinski et al., 2019), but that involved inhibition of other BF neurons in addition to cholinergic neurons. It may be that BF-cholinergic disruption alone can mildly impair complex motor control and increase falls, even if not to a statistically significant extent here. However, the unmasking of concomitant underlying striatal DA deficits with such disruptions of BF-cholinergic activity generates more robust impairments. Indeed, individuals with cholinergic losses and associated cognitive dysfunction such as those with Alzheimer’s disease (Aghourian et al., 2017; Ruberg et al., 1990) exhibit gait impairments and falls but not to the extent as PD patients who fall (Balash et al., 2005; De Melo Coelho et al., 2012).

Traversal times were generally slowed by LED inhibition on both the straight and zig-zag rotating rods, however this effect did not interact significantly with disruption group for any of the conditions. This result corresponds with a previous finding using DREADDs in which CNO administration overall slowed traversal speed, but falls were only elevated by one experimental group (GTs) (Kucinski et al., 2019). This contrasts with prior lesion studies in which dual BF cholinergic and striatal DA lesions caused a differential slowing in rats together with higher falls (Kucinski et al., 2013; Kucinski et al., 2017).

### Falls by sign-trackers as well as goal-trackers

The current results indicate that dual ACh/DA disruption-induced propensity to fall applies equally to females and males, and also equally to sign-trackers (STs) as well as to goal-trackers (GTs). This contrasts to a previous finding that BF disruption increased falls primarily in GTs relative to STs (Kucinski et al., 2019). GTs were previously found also to have higher levels of cholinergic neuromodulation in prefrontal cortex projecting BF neurons (Koshy Cherian et al., 2017) and to perform superiorly on a sustained attention task relative to STs (Kucinski et al., 2022). Thus, disruption of BF activity conceivably has a greater effect on GTs’ more top-down (cognitive) vs bottom-up (habitual) behavioral mode.

However, the previous study which found greater vulnerability to falls in GTs induced by nonspecific chemogenetic BF inhibition did not include striatal DA lesions. 6-OHDA lesions of the dorsomedial striatum may interact with prefrontal systems that update habitual and goal-directed behaviors (Smith & Graybiel, 2013), and impact converging pathways from dorsolateral striatal regions involved in habits and which are affected in PD patients (Redgrave et al., 2010), and which may be relevant to ST performance when combined with BF ACh inhibition. Conversely, the current study explicitly assessed the effects of dual ACh/DA disruption, as well as separate ACh inhibition and 6-OHDA lesions, on vulnerability to falls.

A question may remain as to why GTs were not impaired in the current study by BF ACh inhibition alone? An approach was used that differed from previous studies, which may potentially be related to why it was found that dual but not single system disruption caused elevated falls in both GTs and STs. For example, this study specifically inhibited only ACh neurons in the BF, in contrast to the nonspecific DREADD inhibition of BF neurons used in Kucinski et al., 2019. Nonspecific targeting of the BF with DREADDs may affect other neurotransmitter systems such as GABAeric neurons which could confound the role of the cholinergic system in mediating complex movement (Gritti et al., 2006). Also, CNO injections that work on the time scale of several hours may provide a more tonic inhibition with different dynamics than optogenetic inhibition. The sudden suppression of ACh neuronal activity caused by optogenetic inhibition could conceivably be more disruptive to neuronal computations needed by GTs than the relatively slow and gradual disruption induced by chemogenetic inhibition. It remains possible that GTs are more vulnerable than STs to falls induced by dual disruption that involves more gradual or milder impairment of ACh function than here. But these results suggest that both GT and STs are vulnerable to falls induced by dual ACh/DA disruption. The human population embodies a spectrum of GT and ST behaviors rather than strict categorical divisions into groups, and the higher fall propensity of both groups of rats may more realistically reflect how PD patients are affected by cholinergic system disruption.

### Types of falls

Rats with dual DA and ACh disruption fell more with both 1s LED inhibitory pulses (given twice per 3-meter traversal) and continuous LED inhibition for the duration of traversals. A brief analysis of fall behavior did not reveal differences in types of falls or differences in falls induced by 1s LED inhibition vs continuous LED inhibition. In general, falls with optogenetic disruption appeared to be similar in nature as those observed with chemogenetic disruption (Kucinski et al., 2019). Rats fell both ‘spontaneously’ with no obvious trigger or slowed with low motor vigor and were unable to effectively rebalance and continue forward traversal following slips or missteps, leading to falls.

As in previous MCMCT experiments, females were generally faster to traverse both the straight and zig-zag rotating rods than males, and fell less frequently from the zig-zag rods (Kucinski et al., 2017; Kucinski et al., 2019). This was likely due to males’ larger body size which makes it difficult to navigate tight bends of the zig-zagged sections and re-balance following slips or missteps. Both 1s and continuous LED inhibition increased falls over baseline in males and females on the zig-zag rotating rod in *Dual ACh/DA Disruption* rats, demonstrating that disruption of cholinergic BF neurons impairs complex movement over a range of baseline motor abilities and body sizes.

Histological analysis verified that *Dual ACh/DA Disruption* rats that were more affected by LED inhibition, particularly continuous inhibition for the duration of traversals, had higher expression of the inhibitory optogenetic AAV in the SI and HDB of the BF. This result is consistent with expression of DREADDs that evoked falls with CNO administration (Kucinski et al., 2019). Separately, rats with larger and more specific 6-OHDA lesions of the dorsomedial striatum in the *Dual ACh/DA Disruption* group tended to fall more with continuous LED inhibition, a result that is consistent with previous experiments using the dual system lesions (Kucinski et al., 2013, Kucinski et al., 2015), and similar to human patients (Bohnen et al., 2009). More severe disruptions of either the cholinergic BF or dorsal striatal dopamine system may contribute to increased movement deficits rather than these impairments being a function of disruption of a single system.

Taken together, these results support and extend previous findings in rats with striatal DA losses that cholinergic disruption of BF cortically-projecting neurons precipitates falls and related gait, balance, and postural deficits, similar to human PD patients. These effects were observed consistently in both GT and ST rats with varied cognitive strategies for complex surface traversal. Also, since the optogenetic inhibition reveals the impact of BF cholinergic disruption on falls on a sub-second time scale, therapies that enhance cholinergic output in such a phasic manner may potentially alleviate these dual-system movements deficits, including symptoms of cognitive decline.

## Author contributions

AK conducted the experiments, analyzed the data, and wrote the paper.

## Acknowledgements

The research described in this manuscript was supported by NIH grant P50NS091856 (Morris K. Udall Center for Excellence in Parkinson’s Disease Research). I thank Dr. Kent Berridge and Dr. Roger Albin for comments on drafts of this manuscript and Dr. Martin Sarter (University of Michigan) for input on experimental design.

## References

1. Aghourian, M., Legault-Denis, C., Soucy, J.-P., Rosa-Neto, P., Gauthier, S., Kostikov, A., Gravel, P., & Bédard, M.-A. (2017). Quantification of brain cholinergic denervation in Alzheimer’s disease using PET imaging with [18F]-FEOBV. Molecular Psychiatry, 22(11), 1531–1538. 10.1038/mp.2017.183

2. Albin, R. L., Müller, M. L. T. M., Bohnen, N. I., Spino, C., Sarter, M., Koeppe, R. A., Szpara, A., Kim, K., Lustig, C., & Dauer, W. T. (2021). Α4β2* Nicotinic Cholinergic Receptor Target Engagement in Parkinson Disease GAIT–BALANCE Disorders. Annals of Neurology, 90(1), 130–142. 10.1002/ana.26102

3. Albin, R. L., van der Zee, S., van Laar, T., Sarter, M., Lustig, C., Muller, M. L. T. M., & Bohnen, N. I. (2022). Cholinergic systems, attentional-motor integration, and cognitive control in Parkinson’s disease. Progress in Brain Research, 269(1), 345–371. 10.1016/bs.pbr.2022.01.011

4. Allcock, L. M., Rowan, E. N., Steen, I. N., Wesnes, K., Kenny, R. A., & Burn, D. J. (2009). Impaired attention predicts falling in Parkinson’s disease. Parkinsonism & Related Disorders, 15(2), 110–115. 10.1016/j.parkreldis.2008.03.010

5. Arvidsson, U., Riedl, M., Elde, R., & Meister, B. (1997). Vesicular acetylcholine transporter (Vacht) protein: A novel and unique marker for cholinergic neurons in the central and peripheral nervous systems. The Journal of Comparative Neurology, 378(4), 454–467.

6. Avila, C., Kucinski, A., & Sarter, M. (2020). Complex movement control in a rat model of parkinsonian falls: Bidirectional control by striatal cholinergic interneurons. The Journal of Neuroscience, 40(31), 6049–6067. 10.1523/JNEUROSCI.0220-20.2020

7. Balash, Y., Peretz, C., Leibovich, G., Herman, T., Hausdorff, J. M., & Giladi, N. (2005). Falls in outpatients with Parkinson’s disease: Frequency, impact and identifying factors. Journal of Neurology, 252(11), 1310–1315. 10.1007/s00415-005-0855-3

8. Ballinger, E. C., Ananth, M., Talmage, D. A., & Role, L. W. (2016). Basal forebrain cholinergic circuits and signaling in cognition and cognitive decline. Neuron, 91(6), 1199–1218. 10.1016/j.neuron.2016.09.006

9. Bohnen, N. I., & Albin, R. L. (2009). Cholinergic denervation occurs early in Parkinson disease. Neurology, 73(4), 256–257. 10.1212/WNL.0b013e3181b0bd3d

10. Bohnen, N. I., Yarnall, A. J., Weil, R. S., Moro, E., Moehle, M. S., Borghammer, P., Bedard, M.-A., & Albin, R. L. (2022). Cholinergic system changes in Parkinson’s disease: Emerging therapeutic approaches. The Lancet Neurology, 21(4), 381–392. 10.1016/S1474-4422(21)00377-X

11. Bohnen, N. I., Kaufer, D. I., Hendrickson, R., Ivanco, L. S., Lopresti, B. J., Constantine, G. M., Mathis, Ch. A., Davis, J. G., Moore, R. Y., & DeKosky, S. T. (2006). Cognitive correlates of cortical cholinergic denervation in Parkinson’s disease and parkinsonian dementia. Journal of Neurology, 253(2), 242–247. 10.1007/s00415-005-0971-0

12. Bohnen, N. I., Kaufer, D. I., Ivanco, L. S., Lopresti, B., Koeppe, R. A., Davis, J. G., Mathis, C. A., Moore, R. Y., & DeKosky, S. T. (2003a). Cortical cholinergic function is more severely affected in parkinsonian dementia than in alzheimer disease: An in vivo positron emission tomographic study. Archives of Neurology, 60(12), 1745. 10.1001/archneur.60.12.1745

13. Bohnen, N. I., Kaufer, D. I., Ivanco, L. S., Lopresti, B., Koeppe, R. A., Davis, J. G., Mathis, C. A., Moore, R. Y., & DeKosky, S. T. (2003b). Cortical cholinergic function is more severely affected in parkinsonian dementia than in alzheimer disease: An in vivo positron emission tomographic study. Archives of Neurology, 60(12), 1745. 10.1001/archneur.60.12.1745

14. Bohnen, N. I., Mu ller, M. L. T. M., Koeppe, R. A., Studenski, S. A., Kilbourn, M. A., Frey, K. A., & Albin, R. L. (2009). History of falls in Parkinson disease is associated with reduced cholinergic activity. Neurology, 73(20), 1670–1676. 10.1212/WNL.0b013e3181c1ded6

15. Bohnen, N. I., Muller, M. L. T. M., Kuwabara, H., Cham, R., Constantine, G. M., & Studenski, S. A. (2009). Age-associated striatal dopaminergic denervation and falls in community-dwelling subjects. The Journal of Rehabilitation Research and Development, 46(8), 1045. 10.1682/JRRD.2009.03.0030

16. Breese, G. R., & Traylor, T. D. (1971). Depletion of brain noradrenaline and dopamine by 6-hydroxydopamine. British Journal of Pharmacology, 42(1), 88–99. 10.1111/j.1476-5381.1971.tb07089.x

17. Chen, X.-J., Liu, Y.-H., Xu, N.-L., & Sun, Y.-G. (2021). Multiplexed representation of itch and mechanical and thermal sensation in the primary somatosensory cortex. The Journal of Neuroscience, 41(50), 10330–10340. 10.1523/JNEUROSCI.1445-21.2021

18. d’Angremont, E., Renken, R., Van Der Zee, S., De Vries, E. F. J., Van Laar, T., & Sommer, I. E. C. (2025). Cholinergic denervation patterns in parkinson’s disease associated with cognitive impairment across domains. Human Brain Mapping, 46(2), e70047. 10.1002/hbm.70047

19. Dalrymple, W. A., Huss, D. S., Blair, J., Flanigan, J. L., Patrie, J., Sperling, S. A., Shah, B. B., Harrison, M. B., Druzgal, T. J., & Barrett, M. J. (2021). Cholinergic nucleus 4 atrophy and gait impairment in Parkinson’s disease. Journal of Neurology, 268(1), 95–101. 10.1007/s00415-020-10111-2

20. De Melo Coelho, F. G., Stella, F., De Andrade, L. P., Barbieri, F. A., Santos-Galduróz, R. F., Gobbi, S., Costa, J. L. R., & Gobbi, L. T. B. (2012). Gait and risk of falls associated with frontal cognitive functions at different stages of Alzheimer’s disease. *Aging*, Neuropsychology, and Cognition, 19(5), 644–656. 10.1080/13825585.2012.661398

21. Dimitrov, E. (2024). Optogenetic inhibition of the cortical efferents to the locus ceruleus region of pontine tegmentum causes cognitive deficits. Journal of Integrative Neuroscience, 23(3), 60. 10.31083/j.jin2303060

22. Düzel, S., Münte, T. F., Lindenberger, U., Bunzeck, N., Schütze, H., Heinze, H.-J., & Düzel, E. (2010). Basal forebrain integrity and cognitive memory profile in healthy aging. Brain Research, 1308, 124–136. 10.1016/j.brainres.2009.10.048

23. Gombkoto, P., Gielow, M., Varsanyi, P., Chavez, C., & Zaborszky, L. (2021). Contribution of the basal forebrain to corticocortical network interactions. Brain Structure and Function, 226(6), 1803–1821. 10.1007/s00429-021-02290-z

24. Greenwald, Anthony G., Gonzalez, R., Harris, Richard J., & Guthrie, D. (1996). Effect sizes and p values: What should be reported and what should be replicated? Psychophysiology, 33(2), 175–183. 10.1111/j.1469-8986.1996.tb02121.x

25. Grimbergen, Y. A., Munneke, M., & Bloem, B. R. (2004). Falls in Parkinson’s disease. Current Opinion in Neurology, 17(4), 405–415. 10.1097/01.wco.0000137530.68867.93

26. Gritti, I., Henny, P., Galloni, F., Mainville, L., Mariotti, M., & Jones, B. E. (2006). Stereological estimates of the basal forebrain cell population in the rat, including neurons containing choline acetyltransferase, glutamic acid decarboxylase or phosphate-activated glutaminase and colocalizing vesicular glutamate transporters. Neuroscience, 143(4), 1051–1064. 10.1016/j.neuroscience.2006.09.024

27. Gritton, H. J., Howe, W. M., Mallory, C. S., Hetrick, V. L., Berke, J. D., & Sarter, M. (2016). Cortical cholinergic signaling controls the detection of cues. Proceedings of the National Academy of Sciences, 113(8). 10.1073/pnas.1516134113

28. Grothe, M. J., Labrador-Espinosa, M. A., Jesús, S., Macías-García, D., Adarmes-Gómez, A., Carrillo, F., Camacho, E. I., Franco-Rosado, P., Lora, F. R., Martín-Rodríguez, J. F., Barberá, M. A., Pastor, P., Arroyo, S. E., Vila, B. S., Foraster, A. C., Martínez, J. R., Padilla, F. C., Morlans, M. P., Aramburu, I. G., … Mir, P. (2021). In vivo cholinergic basal forebrain degeneration and cognition in Parkinson’s disease: Imaging results from the COPPADIS study. Parkinsonism & Related Disorders, 88, 68–75. 10.1016/j.parkreldis.2021.05.027

29. Holtzer, R., Friedman, R., Lipton, R. B., Katz, M., Xue, X., & Verghese, J. (2007). The relationship between specific cognitive functions and falls in aging. Neuropsychology, 21(5), 540–548. 10.1037/0894-4105.21.5.540

30. Kehagia, A. A., Barker, R. A., & Robbins, T. W. (2013). Cognitive impairment in parkinson’s disease: The dual syndrome hypothesis. Neurodegenerative Diseases, 11(2), 79–92. 10.1159/000341998

31. Kim, K., Müller, M. L. T. M., Bohnen, N. I., Sarter, M., & Lustig, C. (2019). The cortical cholinergic system contributes to the top-down control of distraction: Evidence from patients with Parkinson’s disease. NeuroImage, 190, 107–117. 10.1016/j.neuroimage.2017.12.012

32. Koshy Cherian, A., Kucinski, A., Pitchers, K., Yegla, B., Parikh, V., Kim, Y., Valuskova, P., Gurnani, S., Lindsley, C. W., Blakely, R. D., & Sarter, M. (2017). Unresponsive choline transporter as a trait neuromarker and a causal mediator of bottom-up attentional biases. The Journal of Neuroscience, 37(11), 2947–2959. 10.1523/JNEUROSCI.3499-16.2017

33. Kucinski, A., Albin, R. L., Lustig, C., & Sarter, M. (2015). Modeling falls in Parkinson’s disease: Slow gait, freezing episodes and falls in rats with extensive striatal dopamine loss. Behavioural Brain Research, 282, 155–164. 10.1016/j.bbr.2015.01.012

34. Kucinski, A., Avila, C., & Sarter, M. (2022). Basal forebrain chemogenetic inhibition converts the attentional control mode of goal-trackers to that of sign-trackers. Eneuro, 9(6), ENEURO.0418-22.2022. 10.1523/ENEURO.0418-22.2022

35. Kucinski, A., De Jong, I. E. M., & Sarter, M. (2017). Reducing falls in Parkinson’s disease: Interactions between donepezil and the 5-HT_6_ receptor antagonist idalopirdine on falls in a rat model of impaired cognitive control of complex movements. European Journal of Neuroscience, 45(2), 217–231. 10.1111/ejn.13354

36. Kucinski, A., Kim, Y., & Sarter, M. (2019). Basal forebrain chemogenetic inhibition disrupts the superior complex movement control of goal-tracking rats. Behavioral Neuroscience, 133(1), 121–134. 10.1037/bne0000290

37. Kucinski, A., Lustig, C., & Sarter, M. (2018). Addiction vulnerability trait impacts complex movement control: Evidence from sign-trackers. Behavioural Brain Research, 350, 139–148. 10.1016/j.bbr.2018.04.045

38. Kucinski, A., Paolone, G., Bradshaw, M., Albin, R. L., & Sarter, M. (2013). Modeling fall propensity in Parkinson’s disease: Deficits in the attentional control of complex movements in rats with cortical-cholinergic and striatal–dopaminergic deafferentation. The Journal of Neuroscience, 33(42), 16522–16539. 10.1523/JNEUROSCI.2545-13.2013

39. Kucinski, A., & Sarter, M. (2021). Reduction of falls in a rat model of PD falls by the M1 PAM TAK-071. Psychopharmacology, 238(7), 1953–1964. 10.1007/s00213-021-05822-x

40. LaPointe, L. L., Stierwalt, J. A. G., & Maitland, C. G. (2010). Talking while walking: Cognitive loading and injurious falls in Parkinson’s disease. International Journal of Speech-Language Pathology, 12(5), 455–459. 10.3109/17549507.2010.486446

41. Liu, A. K. L., Chang, R. C.-C., Pearce, R. K. B., & Gentleman, S. M. (2015). Nucleus basalis of Meynert revisited: Anatomy, history and differential involvement in Alzheimer’s and Parkinson’s disease. Acta Neuropathologica, 129(4), 527–540. 10.1007/s00401-015-1392-5

42. Ma, M., & Luo, M. (2012). Optogenetic activation of basal forebrain cholinergic neurons modulates neuronal excitability and sensory responses in the main olfactory bulb. The Journal of Neuroscience, 32(30), 10105–10116. 10.1523/JNEUROSCI.0058-12.2012

43. Mailly, P., Aliane, V., Groenewegen, H. J., Haber, S. N., & Deniau, J.-M. (2013). The rat prefrontostriatal system analyzed in 3d: Evidence for multiple interacting functional units. The Journal of Neuroscience, 33(13), 5718–5727. 10.1523/JNEUROSCI.5248-12.2013

44. Meyer, P. J., Lovic, V., Saunders, B. T., Yager, L. M., Flagel, S. B., Morrow, J. D., & Robinson, T. E. (2012). Quantifying individual variation in the propensity to attribute incentive salience to reward cues. PLoS ONE, 7(6), e38987. 10.1371/journal.pone.0038987

45. Norman, K. J., Riceberg, J. S., Koike, H., Bateh, J., McCraney, S. E., Caro, K., Kato, D., Liang, A., Yamamuro, K., Flanigan, M. E., Kam, K., Falk, E. N., Brady, D. M., Cho, C., Sadahiro, M., Yoshitake, K., Maccario, P., Demars, M. P., Waltrip, L., … Morishita, H. (2021). Post-error recruitment of frontal sensory cortical projections promotes attention in mice. Neuron, 109(7), 1202–1213.e5. 10.1016/j.neuron.2021.02.001

46. Paolone, G., Angelakos, C. C., Meyer, P. J., Robinson, T. E., & Sarter, M. (2013). Cholinergic control over attention in rats prone to attribute incentive salience to reward cues. Journal of Neuroscience, 33(19), 8321–8335. 10.1523/JNEUROSCI.0709-13.2013

47. Pitchers, K. K., Kane, L. F., Kim, Y., Robinson, T. E., & Sarter, M. (2017). ‘Hot’ vs. ‘cold’ behavioural-cognitive styles: Motivational-dopaminergic vs. cognitive-cholinergic processing of a Pavlovian cocaine cue in sign- and goal-tracking rats. European Journal of Neuroscience, 46(11), 2768–2781. 10.1111/ejn.13741

48. Pitchers, K. K., Phillips, K. B., Jones, J. L., Robinson, T. E., & Sarter, M. (2017). Diverse roads to relapse: A discriminative cue signaling cocaine availability is more effective in renewing cocaine seeking in goal trackers than sign trackers and depends on basal forebrain cholinergic activity. Journal of Neuroscience, 37(30), 7198–7208. 10.1523/JNEUROSCI.0990-17.2017

49. Quik, M., Boyd, J. T., Bordia, T., & Perez, X. (2019). Potential therapeutic application for nicotinic receptor drugs in movement disorders. Nicotine & Tobacco Research, 21(3), 357–369. 10.1093/ntr/nty063

50. Redgrave, P., Rodriguez, M., Smith, Y., Rodriguez-Oroz, M. C., Lehericy, S., Bergman, H., Agid, Y., DeLong, M. R., & Obeso, J. A. (2010). Goal-directed and habitual control in the basal ganglia: Implications for Parkinson’s disease. Nature Reviews Neuroscience, 11(11), 760–772. 10.1038/nrn2915

51. Robinson, T. E., Yager, L. M., Cogan, E. S., & Saunders, B. T. (2014). On the motivational properties of reward cues: Individual differences. Neuropharmacology, 76, 450–459. 10.1016/j.neuropharm.2013.05.040

52. Rochester, L., Yarnall, A. J., Baker, M. R., David, R. V., Lord, S., Galna, B., & Burn, D. J. (2012). Cholinergic dysfunction contributes to gait disturbance in early Parkinson’s disease. Brain, 135(9), 2779–2788. 10.1093/brain/aws207

53. Ruberg, M., Mayo, W., Brice, A., Duyckaerts, C., Hauw, J. J., Simon, H., LeMoal, M., & Agid, Y. (1990). Choline acetyltransferase activity and [ 3H]vesamicol binding in the temporal cortex of patients with Alzheimer’s disease, Parkinson’s disease, and rats with basal forebrain lesions. Neuroscience, 35(2), 327–333. 10.1016/0306-4522(90)90086-J

54. Saito, Y., Osako, Y., Odagawa, M., Oisi, Y., Matsubara, C., Kato, S., Kobayashi, K., Morita, M., Johansen, J. P., & Murayama, M. (2025). Amygdalo-cortical dialogue underlies memory enhancement by emotional association. Neuron, 113(6), 931–948.e7. 10.1016/j.neuron.2025.01.001

55. Sarter, M. (1998). Age-related changes in rodent cortical acetylcholine and cognition: Main effects of age versus age as an intervening variable. Brain Research Reviews, 27(2), 143–156. 10.1016/S0165-0173(98)00003-4

56. Sarter, M., Albin, R. L., Kucinski, A., & Lustig, C. (2014). Where attention falls: Increased risk of falls from the converging impact of cortical cholinergic and midbrain dopamine loss on striatal function. Experimental Neurology, 257, 120–129. 10.1016/j.expneurol.2014.04.032

57. Sarter, M., & Lustig, C. (2020). Forebrain cholinergic signaling: Wired and phasic, not tonic, and causing behavior. The Journal of Neuroscience, 40(4), 712–719. 10.1523/JNEUROSCI.1305-19.2019

58. Sarter, M., & Paolone, G. (2011). Deficits in attentional control: Cholinergic mechanisms and circuitry-based treatment approaches. Behavioral Neuroscience, 125(6), 825–835. 10.1037/a0026227

59. Schliebs, R., & Arendt, T. (2011). The cholinergic system in aging and neuronal degeneration. Behavioural Brain Research, 221(2), 555–563. 10.1016/j.bbr.2010.11.058

60. Schulz, J., Pagano, G., Fernández Bonfante, J. A., Wilson, H., & Politis, M. (2018). Nucleus basalis of Meynert degeneration precedes and predicts cognitive impairment in Parkinson’s disease. Brain, 141(5), 1501–1516. 10.1093/brain/awy072

61. Smith, K. S., & Graybiel, A. M. (2013). A dual operator view of habitual behavior reflecting cortical and striatal dynamics. Neuron, 79(2), 361–374. 10.1016/j.neuron.2013.05.038

62. St. Peters, M., Demeter, E., Lustig, C., Bruno, J. P., & Sarter, M. (2011). Enhanced control of attention by stimulating mesolimbic-corticopetal cholinergic circuitry. Journal of Neuroscience, 31(26), 9760–9771. 10.1523/JNEUROSCI.1902-11.2011

63. Stuart, S., Morris, R., Giritharan, A., Quinn, J., Nutt, J. G., & Mancini, M. (2020). Prefrontal cortex activity and gait in parkinson’s disease with cholinergic and dopaminergic therapy. Movement Disorders, 35(11), 2019–2027. 10.1002/mds.28214

64. Wolf, D., Grothe, M., Fischer, F. U., Heinsen, H., Kilimann, I., Teipel, S., & Fellgiebel, A. (2014). Association of basal forebrain volumes and cognition in normal aging. Neuropsychologia, 53, 54–63. 10.1016/j.neuropsychologia.2013.11.002

65. Woollacott, M., & Shumway-Cook, A. (2002). Attention and the control of posture and gait: A review of an emerging area of research. Gait & Posture, 16(1), 1–14. 10.1016/S0966-6362(01)00156-4

66. Yager, L. M., Pitchers, K. K., Flagel, S. B., & Robinson, T. E. (2015). Individual variation in the motivational and neurobiological effects of an opioid cue. Neuropsychopharmacology, 40(5), 1269–1277. 10.1038/npp.2014.314

67. Yager, L. M., & Robinson, T. E. (2013). A classically conditioned cocaine cue acquires greater control over motivated behavior in rats prone to attribute incentive salience to a food cue. Psychopharmacology, 226(2), 217–228. 10.1007/s00213-012-2890-y

68. Yarnall, A., Rochester, L., & Burn, D. J. (2011). The interplay of cholinergic function, attention, and falls in Parkinson’s disease. Movement Disorders, 26(14), 2496–2503. 10.1002/mds.23932

69. Záborszky, L., Gombkoto, P., Varsanyi, P., Gielow, M. R., Poe, G., Role, L. W., Ananth, M., Rajebhosale, P., Talmage, D. A., Hasselmo, M. E., Dannenberg, H., Minces, V. H., & Chiba, A. A. (2018). Specific basal forebrain–cortical cholinergic circuits coordinate cognitive operations. The Journal of Neuroscience, 38(44), 9446–9458. 10.1523/JNEUROSCI.1676-18.2018

70. Zaborszky, L., Hoemke, L., Mohlberg, H., Schleicher, A., Amunts, K., & Zilles, K. (2008). Stereotaxic probabilistic maps of the magnocellular cell groups in human basal forebrain. NeuroImage, 42(3), 1127–1141. 10.1016/j.neuroimage.2008.05.055

71. Zant, J. C., Kim, T., Prokai, L., Szarka, S., McNally, J., McKenna, J. T., Shukla, C., Yang, C., Kalinchuk, A. V., McCarley, R. W., Brown, R. E., & Basheer, R. (2016). Cholinergic neurons in the basal forebrain promote wakefulness by actions on neighboring non-cholinergic neurons: An opto-dialysis study. The Journal of Neuroscience, 36(6), 2057–2067. 10.1523/JNEUROSCI.3318-15.2016

